# Probing tau citrullination in Alzheimer’s disease brains and mouse models of tauopathy

**DOI:** 10.1101/2024.07.06.601399

**Authors:** Huimin Liang, Jerry B. Hunt, Chao Ma, Andrii Kovalenko, John Calahatian, Cecelie Pedersen, Haiying Lui, Junyan Li, Malina Serrano, Danielle Blazier, Mallory Watler, Patricia Rocha-Rangel, Christopher Saunders, Laura J. Blair, Leonid Breydo, Kevin Nash, Zainuddin Quadri, Brian Kraemer, Peter Nelson, Christopher Norris, Erin L. Abner, Vladimir N. Uversky, Dale Chaput, Maj-Linda B. Selenica, Daniel C. Lee

## Abstract

Tauopathies, which include Alzheimer’s disease (AD) share a common defining factor, namely misfolded tau protein. However, the “upstream” etiology and downstream clinical manifestations of tauopathies are quite diverse. Tau deposition elicits different pathological phenotypes and outcomes depending on the tau strain and regional susceptibility. Posttranslational modifications (PTM) can alter tau structure, function, networks, and its pathological sequalae. We uncovered a novel PTM of tau, named citrullination, caused by peptidyl arginine deiminase (PAD) enzymes. PAD induced citrullination irreversibly converts arginine residues to citrulline, producing net loss of positive charge, elimination of pi-pi interactions, and increased hydrophobicity. We observed increased PAD2 and PAD4 in Alzheimer’s disease (AD) brain and that they both can citrullinate tau. Tau can become citrullinated by PADs at all 14 arginine residues throughout the N-terminal domain (N-term), proline-rich domain (PR), microtubule binding repeat domain (MBR), and C-terminal domain (C-term) on full length tau (2N4R). Citrullination of tau impacts fibrillization and oligomerization rates in aggregation assays. Utilizing a panel of novel citrullinated tau (citR tau) antibodies, we identified citrullination of tau *in vitro*, several animal models of tauopathies, and Alzheimer’s disease (AD). CitR tau increased with Braak stage and was enriched in AD brains with higher phospho-tau burden. This work provides a new area of tau biology that signifies further consideration in the emerging spectrum of tauopathies and its clinical understanding.

## Introduction

Tauopathies including Alzheimer’s disease (AD) cause a devastating group of disorders that track with the tau deposition. Decades of research conclude that hallmarks of tauopathies consist of loss of tau function, transition from soluble to insoluble tau aggregates and neurofibrillary tangles (NFT), neurodegeneration, and ultimately cognitive impairment. Electron Cryo-microscopy (CryoEm) has uncovered the diversity of tau proteoforms for the evolving class of tauopathies. Increasing reports illustrate how tau dysfunction impacts critical physiological processes, including mitochondria bioenergetics and synaptic plasticity(Tracy, Madero-Perez et al. 2022), and autophagy impairment (Dickey, Kraft et al. 2009, Dolan and Johnson 2010, Cohen, Guo et al. 2011), that subsequently lead to different clinical syndromes. There is an urgent need to understand how different variants, strains, and proteoforms of tau influence biology for anti-tau approaches.

The mammalian genome codes for only 20 natural amino acids, but diversity of posttranslational modifications (PTMs) provide more flexibility in protein structure and function that can produce more than one hundred secondary (derived) amino acids (Uy and Wold 1977), thus allowing unique and new features for a single protein. Tau protein is a substrate for several PTMs including hyperphosphorylation, acetylation, ubiquitination, methylation, glycosylation, nitration, and others less well studied (Cohen, Guo et al. 2011, Morris, Knudsen et al. 2015, Alquezar, Arya et al. 2020). Phosphorylation is recognized as the most studied tau PTM and remains the best understood, where the hyperphosphorylation is thought to initiate the pathogenic sequalae of tau aggregation (Lee, Goedert et al. 2001, Fontaine, Sabbagh et al. 2015, Simic, Babic Leko et al. 2016). Recent studies using cryoEM technology have been instrumental in investigating the role of PTMs in promoting strain and conformation differences, (Arakhamia, Lee et al. 2020) signifying a need to understand other PTMs in tauopathies. Given the dynamic complexity of PTMs in tauopathies (Carlomagno, Chung et al. 2017, Ercan-Herbst, Ehrig et al. 2019, Schaffert and Carter 2020), it is plausible that less well-studied modifications add to tau diversity and contribute to the clinical spectrum.

Citrullination (citR; also called deamination) is an irreversible PTM catalyzed by peptidyl arginine deiminases (PADs) that involves the conversion of the amino acid arginine to citrulline within proteins and peptides (Anzilotti, Pratesi et al. 2010). Citrullination yields the loss of a positive charge coupled with a 0.98 Da increase in mass (Rohrbach, Slade et al. 2012), which increases hydrophobicity of the structural moiety (De Ceuleneer, Van Steendam et al. 2012). In addition, neutralization of positively charged arginines significantly alter protein structure as well as stabilization of protein-protein, protein-DNA, protein-RNA, interactions (Gyorgy, Toth et al. 2006, Wong, Demers et al. 2015). Peptidyl arginine deiminase (*PADI)* genes, encoding the 5 human PAD isoforms (PAD1-5), are located in a single gene cluster on chromosome 1p36.1 in humans and chromosome 4pE1 in mice (Vossenaar, Zendman et al. 2003, Wang, Chen et al. 2022). Influx of, or increased, intracellular calcium activates PADs in a coordinated manner through five calcium binding sites (C1-C5) to form a catalytically competent active site (Arita, Hashimoto et al. 2004, Rohrbach, Slade et al. 2012, Mondal and Thompson 2019). Reports illustrate increased PAD4 expression throughout the cytoplasm of pyramidal neurons with age and in AD human brain (Ishigami, Ohsawa et al. 2005, Wang, Chen et al. 2022). In addition, previous studies demonstrated PAD2 and PAD4 co-localization with citrullinated proteins in the cortex and hippocampus of AD brain tissue (Ishigami, Ohsawa et al. 2005, Acharya, Nagele et al. 2012). Acharya and colleagues provided further evidence of co-localization of pan-citrullinated proteins with amyloid beta and PAD4 in the cytoplasm of cortical pyramidal and hippocampal hilus neurons (Acharya, Nagele et al. 2012). Finally, several citrullinated proteins were identified in various regions of AD brain tissue including GFAP, MBP, HSP90, ENO1, NSE, and PTCD2. Additionally, amyloid beta can also become citrullinated during AD (Ishigami, Ohsawa et al. 2005, Mukherjee, Perez et al. 2021).

Herein, we show that full-length tau is a client of both PAD4 and PAD2, which can become fully citrullinated at all 14 arginine residues using recombinant protein systems. Development of a panel of highly specific citR tau antibodies further demonstrated the extent of tau citrullination in cellular and animal models of tauopathy as well as human AD brain tissue. In tauopathy mice with FTD mutations, PAD4 increased early during tau pathogenesis along with citR tau, which accumulated in soluble and detergent soluble fractions. CitR tau was also increased in human AD tissue with higher phospho tau burden. Overall, this study provides a necessary framework to understand the role of citrullination as a common PTM in AD and tauopathies.

## Material and Methods

### Recombinant tau and *in vitro* citrullination

Recombinant human 2N4R (441 tau protein) was purchased by (cat#T1001, rPeptide, GA, USA). For in vitro citrullination, 2 μg (1 μg/ μl) of recombinant tau was incubated with 2 µg (1 μg / μl) or decreasing concentrations of Peptidyl Arginine Deaminases (PAD2 or PAD4, MI, USA) (Cayman, MI.USA) in PAD reaction buffer (50mM Boric Acid, 2-10mM CaCl2 Dihydrous, pH 8.0, Cayman, 10500, GA, USA) (Cayman, MI.USA) at 37°C for 90 minutes.

### Mass Spectroscopy

Samples from enzyme reactions with tau protein incubated with PAD2 or PAD4 were separated by SDS-PAGE and Coomassie-stained for visualization. Bands were excised from the gel and processed for LC-MS/MS analysis. Briefly, each gel piece was minced and de-stained before being reduced with dithiothreitol (DTT), alkylated with iodoacetamide (IAA), and finally digested with Trypsin/Lys-C overnight at 37°C. Peptides were extracted using 50/50 acetonitrile (ACN)/H_2_O/0.1% formic acid and dried in a vacuum concentrator. Peptides were resuspended in 98%H_2_O/2%ACN/0.1% formic acid for LC-MS/ MS analysis.

Peptides were separated on a 50 cm C18 reversed-phase UHPLC column (Thermo Fisher Scientific, MA, USA) using an Ultimate3000 UHPLC (Thermo Fisher Scientific, MA, USA) with a 60 minute gradient (4-40%ACN/0.1% formic acid) and analyzed on a hybrid quadrupole-Orbitrap mass spectrometer (Q Exactive Plus, Thermo Fisher Scientific, MA, USA) using data-dependent acquisition in which the top 10 most abundant ions are selected for MS/MS analysis. Full MS survey scans were acquired at 70,000 resolution.

Raw data files were processed in MaxQuant (www.maxquant.org) and searched against the current Uniprot *Homo sapiens* protein sequence database. Search parameters included constant modification of cysteine by carbamidomethylation, and the variable modifications, methionine oxidation, arginine citrullination, as well as asparagine and glutamine deamidation. Proteins are identified using the filtering criteria of 1% protein and peptide false discovery rate, with a peptide tolerance of 4.5ppm.

To ensure the correct identification of the site-specific citrullination of Tau411, several factors were taken into consideration. MS/MS spectra were manually annotated to verify the sequence coverage of each citrulline containing peptide. Verification process further included consideration of deamidation of asparagine and glutamine, which results in an identical mass shift as the citrullination of arginine. While deamidation cannot be differentiated from citrullination based on mass, deamidation does not affect the charge of the amino acid residue (under acidic LC-MS conditions), however citrulline neutralizes a positively charged arginine. The reduced positive charge/increased hydrophobicity of a peptide containing a citrullinated arginine results in later elution time compared to its unmodified counterpart (Raijmakers, van Beers et al. 2012). In addition to manual inspection of the MS/MS spectra, we took into consideration peptide retention times. For this, the extracted ion chromatogram (XIC) for each peptide identified showed a later elution time for citrullinated peptides compared with the unmodified version. Citrullination was only considered if they could be reliably distinguished from asparagine or glutamine deamidation, and if there was sufficient b- and y-fragment ion coverage surrounding the modified site.

### Generation of polyclonal citR tau antibodies

The rabbit polyclonal antibodies raised against citrullinated tau at arginine epitopes R5, R23, R155, R170, R194, R209, R221, R230, R349, R379 and R406 was produced (21^st^ Century Biochemicals) using their custom polyclonal antibody cross-affinity purification service. Citrullinated specific antibodies were purified using affinity chromatography with modified and unmodified peptides and Q/C immunodepleted. The initial validation for citR-specificity of cross-affinity purified antibodies was performed in recombinant protein, preabsorbed citR tau peptide controls, transgenic tau mouse model and AD human brain tissue.

### Cell culture and treatment

The iHEK-P301L cells (iHEK cells, generously gifted by late Dr. Chad Dickey at University of South Florida, FL, USA) were cultured in DMEM supplemented with 10% FBS (Thermo Fisher Scientific, MA, USA) and 1% penicillin/streptomycin (Corning, NY, USA). Blasticidin (3 μg/ml, Thermo Fisher Scientific, MA, USA) and Zeocin (125 μg/ml, Thermo Fisher Scientific, MA, USA) were added in cell media. Inducible cells were incubated with 1 μg of tetracycline (Sigma Aldrich, MO, USA) and transfected at 70% confluence with PE-GFP or pCMV6-PAD4 constructs (Origene, MR216758, MD, US) for 72 hrs. Transfections were performed with 2.5 μL of Lipofectamine 2000 (Thermo Fisher Scientific, MA, USA) per 1 μg of DNA, which was incubated in serum-free Opti-MEM for 5 min before adding to the iHEK cells. Post-transfection, cells were washed and harvested with PBS and immediately followed by lysis in radio-immunoprecipitation assay buffer (RIPA, 50 mM Tris HCl, 150 mM NaCl, 2 mM EDTA, 0.1% SDS, 0.5% sodium deoxycholate, and 1% Triton x-100, pH 7.4), including protease inhibitor (Sigma Aldrich, MO, USA), phosphatase inhibitor cocktail 2 and 3 purchased from Sigma Aldrich, MO, US, phenylmethylsulfonyl fluoride (0.1 M PMSF, Acros Organics, NJ, USA) and DNase (5 mg/mL, Alfa Aesar, MA, USA) each at a 1:100 dilution. Whole cell fraction is analyzed by western blot.

### Animal models

All animal testing procedures were approved by the Institutional Animal Care and Use Committee of the University of South Florida and University of Kentucky and were performed in accordance with the eighth edition of the “Guide for the Care and Use of Laboratory Animals,” published by the National Academy of Science, the National Academies Press, Washington, DC (2011).

The regulatable Tg4510 (rTg4510) mouse and littermates at 4-, 8-, 13-, and 16-months years old were used in this study (n=5-8 mice/genotype). The rTg4510 mice harbors the *MAPT P301L* mutation (*tetO-MAPT*P301L*), which is associated with an autosomal dominantly inherited dementia referred to as frontotemporal dementia and parkinsonism linked to chromosome 17 (FTDP-17) (Hutton, Lendon et al. 1998). Parental mutant tau and tetracycline-controlled transactivator (tTA) protein mouse lines were maintained separately and bred to produce rTg4510, non-transgenic and tTA only littermates as described previously (Santacruz, Lewis et al. 2005). The 9-month-old PS19 mice (human *MAPT P301S, 1N4R; prion promoter*) and non-transgenic littermates (N=3-4 mice/genotype). The PS19 mice exhibit tau phosphorylation and pre-tangle pathology at 4-6 month of age while developing tau deposition, NFTs, oligomeric tau, neuron loss, regional atrophy, inflammation, and cognitive deficits with later age (Yoshiyama, Higuchi et al. 2007). Finally, 9-month-old 4RTg2652 mice (human *MAPT, 1N4R; Thy1 promoter*) expressing human wildtype tau (Wheeler, McMillan et al. 2015) and non-transgenic littermates were also included (N=3-4 mice/genotype). The 4RTg2652 mice develop early non-progressive tau, pre-tangle phosphorylated tau, sparse oligomeric species, but undetectable fibrillar tau pathology.

### Human post-mortem samples

Human post-mortem tissues were obtained from the National Institute of Health (NIH) NeuroBio Bank and de-identified according to institutional bioethics guidelines. Consent for autopsy was obtained from probands or their legal representative in accordance with NIH institutional review boards. Tissue from a total of 25 cases were selected based on NIH diagnosis of AD neuropathological change (Montine, Phelps et al. 2012) (N=15; age range 67-100; [81.8 ± 1.8, mean ± S.E.M]) or neurologically healthy cases (N=10; age range 69-89; [78.5 ± 2.4, mean ± S.E.M]). The average PMI was (AD=14.9 ± 4.2 hours, mean ± SD; Controls=16.1 ± 4.4 hours, mean ± SD) (Supplementary Table 1, 2). Samples were characterized by sex, Braak Neurofibrillary Tangle Staging, and phospho-tau using AT8 antibody immunohistochemistry (Thermo Fisher Scientific, MA, USA).

### Protein extraction and western blot

Brain cortical tissue were homogenized in Buffer A (10mM Tris-HCl, 80mM NaCl, 1mM MgCl_2_, 1mM EGTA, 0.1mM EDTA, 100mM DTT, 1mM PMSF, protease inhibitor cocktail, and phosphatase inhibitor cocktails II and III, (Sigma Aldrich, MO, USA) and ultracentrifuged at 150,000 x g for 70min at 4°C. Supernatant (S1) was kept as “soluble fraction.” The pellet (P1) was resuspended in Buffer B (10mM Tris-HCl, 850mM NaCl, 1Mm EGTA, 10% sucrose, (Sigma Aldrich, MO, USA) and centrifuged at 14,000 x g for 10min at 4°C to remove debris. Supernatant was incubated with 1% sarkosyl for 1h at room temperature. After incubation, the sample was ultracentrifuged at 150,000 x g for 40min at 4°C. The supernatant (S2) was collected as “sarkosyl-soluble fraction”. The pellet (P3) was resuspended in ice-cold PBS and sonicated as the “sarkosyl-insoluble fraction”. Protein concentrations were determined by the BCA protein assay kit (Pierce, IL, USA). Equal amounts of proteins (5μg/well for soluble fraction, 1μg/well for insoluble fraction) were loaded in each well of a 4–12% Bis-tris gels and transferred to a 0.2 μm pore size nitrocellulose membrane and immunoblotted with Tau CitR antibodies (rabbit polyclonal, anti-citR5, anti-citR23, anti-citR155, anti-citR170, anti-citR194, anti-citR209, anti-citR221, anti-citR230, anti-citR349, anti-citR379, anti-citR406; 21^st^ Century Biochemicals Inc, MA, USA), PAD2 and 4 rabbit polyclonal antibodies (ProteinTech, cat# 12110-1-AP and cat# 17373-1-AP IL, USA), total tau (Agilent Technologies, A002401-2, CA, USA), PHF1 (kindly gift from late Dr. P. Davis), AT180, pSer199/202 tau (Anaspec, cat# 54963-025 CA, USA), pS214 (Abcam, cat# ab170892 Cambridge, UK) and anti-tau phosphorylated at Ser202/Thr205 (AT8) (Thermo Fisher Scientific, MN1020, MA,USA) at 1:1000-fold dilution. Blots were developed using ECL (Pierce, cat # 32106, IL, USA) on Azure 600 imager (AzureBiosystems, CA, USA). Band intensities were quantified by densitometric analysis using AlphaEase software (Alpha Innoch, CA, USA) and normalized to actin band intensity (soluble fraction) or total protein (insoluble fraction). Note: All western blots were run on separate membranes and were only re-probed for the housekeeping protein actin or GAPDH. Thus, all western blot images in figures represent the original immunoblots.

### Tissue collection and Immunohistochemistry

Mice from different ages were weighed, overdosed with a euthanizing solution containing pentobarbital and perfused with 25 mL of 0.9% saline solution. Brains were collected following saline perfusion and were hemisected down the sagittal midline. One hemisphere was dissected and frozen on dry ice for biochemical studies. The second hemisphere was immersion-fixed in 4% paraformaldehyde for 24 hours and cryoprotected in successive incubations of 10%, 20%, and 30% solutions of sucrose for 24 hours in each solution. Subsequently, the fixed hemispheres were frozen on a cold stage and sectioned in the horizontal plane (25 µm thickness) using a sliding microtome. Brain sections were stored in Dulbecco’s phosphate-buffered saline (DPBS, Thermo Fisher Scientific, MA, USA) with 10 mM sodium azide (Thermo Fisher Scientific, MA, USA) solution at 4 °C for immunohistochemistry.

Stereological principles were employed to select the sections stained for each marker. Immunohistochemical procedural methods were previously described (Gordon, Holcomb et al. 2002). Six to eight sections from each animal or two sections/ human brain tissue were placed in a multi-sample staining tray and endogenous peroxidase was blocked (10% methanol, 3% H_2_O_2_ in phosphate buffered saline, 10 mM NaPO_4_, 0.8%NaCl, pH 7.4, PBS; 30 min, (Thermo Fisher Scientific, MA.USA). Tissue was permeabilized with 0.2% lysine, 1% Triton X-100 (Thermo Fisher Scientific, MA.USA) in PBS solution and incubated overnight in primary antibody. The following primary antibodies were used for immunohistochemistry: Tau citR antibodies as described above (21^st^ Century Biochemicals Inc, MA, USA), HT7 (ThermoFisher Scientific, cat# MN1000 MA, USA), PAD2 and 4 antibodies (ProteinTech, IL, USA). Sections were washed in PBS, then incubated in corresponding biotinylated secondary antibody (Vector Laboratories, CA, USA). For colorimetric analysis the tissue was washed after 2 hours and incubated with Vectastain® Elite® ABC kit (Vector Laboratories, CA, USA) for enzyme conjugation. Finally, sections were stained using 0.05% diaminobenzidine, 0.5% nickel ammonium sulfate and 0.03% H_2_O_2_. Tissue sections were dehydrated and permanently mounted onto slides with a cover slip and prepared for analysis.

For immunofluorescence, sections were washed in PBS after primary antibody incubation and incubated with goat anti-mouse Alexa-488 (ThermoFisher Scientific, cat# A-10680) or goat anti-rabbit 594 (ThermoFisher Scientific, cat# A-11012) with prolong gold with DAPI (ThermoFisher Scientific, cat# P36941).

For colorimetric analysis the tissue was washed after 2 hours and incubated with Vectastain® Elite® ABC kit (Vector Laboratories, CA, USA) for enzyme conjugation. Finally, sections were stained using 0.05% diaminobenzidine, 0.5% nickel ammonium sulfate and 0.03% H_2_O_2_. Tissue sections were dehydrated and permanently mounted onto slides with a cover slip and prepared for analysis. For each immunohistochemical assay, 6-8 sections were omitted from primary antibody incubation to evaluate the nonspecific reaction of the secondary antibody.

### Histology

Gallyas staining was performed as described (Lee, Rizer et al. 2010, Hunt, Nash et al. 2015, Joly-Amado, Hunter et al. 2020). Briefly, slides were treated with 5% periodic acid (Thermo Fisher Scientific, MA, USA) for 5 min, washed with water, and incubated sequentially for 1 min in silver iodide (Thermo Fisher Scientific, MA, USA) and 10 min with 0.5% acetic acid (Thermo Fisher Scientific, MA, USA) solutions prior to being placed in developer solution (2.5% sodium carbonate, 0.1% ammonium nitrate, 0.1% silver nitrate, 1% tungstosilicic acid, 0.7% formaldehyde (Thermo Fisher Scientific, MA, USA). Slides were treated with 0.5% acetic acid to stop the reaction, then incubated with 0.1% gold chloride (Thermo Fisher Scientific, MA, USA) and placed in 1% sodium thiosulfate (Thermo Fisher Scientific, MA.USA). Following a final wash in water, slides were rinsed through 3 changes of 100% ethanol, cleared through 3 changes of xylene, and mounted with a coverslip via Di-N-butyl phthalate in xylene (DPX; VWR, PA, USA).

### Tissue Imaging and quantification

Immuno-labeled sections were imaged using a Zeiss Axio Scan Z1 digital slide scanner at various magnification as indicated in each figure legend. Digital images of each slide and its sections were analyzed for threshold-defined pixel-positive area fraction using custom-designed image analysis software (Nearcyte, Zeiss, PA, US). Values for all sections from the same mouse/ human subjects were averaged to represent a single value for that region in subsequent statistical analysis. For analyses, an investigator blinded to the animal genotypes or the human disease stage captured the regional tau pathology and citrullination.

### Statistical analyses

Two-tailed student’s t-test, one-way analysis of variance (ANOVA) with Fisher’s LSD post hoc analysis, or two-way ANOVA with pairwise comparison between genotype and age were used as detailed in the figure legends. Values were considered significant if p < 0.05. Graphs were generated using GraphPad Prism 9.0 analysis software. For western blots, graphs that lack error bars from the control values represent independent experiments run separately and normalized to a value of 1. N-value in this study is depicted as the number of animals / human samples and indicated in the figure legends.

## Results

### PAD expression is increased during tauopathy in AD human tissue

Several reports show increased PAD2 and PAD4 in AD brain, therefore we measured phospho tau expression, PAD2 and PAD4 proteins in AD post-mortem cortex obtained from the NIH NeuroBio Tissue bank. A total of 25 cases diagnosed based on NIH criteria for AD (N=15, F/M 7/8) or neurologically healthy cases (N=10, F/M 5/5), and designated by Braak Stage (**Table 1**). We performed immunohistochemistry and western blot analysis on these cases. First, we measure total tau levels (HT7), tau AT8 and PHF1 in control tissue (Braak II and Braak IV) and AD tissue (Braak VI). Total tau was observed in the case with Braak IV, however AT8 or PHF1 did not stain control tissue Braak II and IV (**Figure 1A**). Conversely, total tau, AT8, and PHF1 readily labeled tau in several cases with AD braak VI. We next performed western blot analysis on tau epitopes for several cases diagnosed with AD but not pathologically confirmed or no information provided. We found that several cases showed lower AT8/ PHF1 expression, which were segregated into AD (+)/ AT8 (−) or AD (+)/ AT8 (+). Tau PHF1, AT180, pSer199/202, increased in the AD (+)/ AT8 (+) (**Figure 1 B, E**), whereas tau pSer214 increased in both AD (+)/ AT8 (−) or AD (+)/ AT8 (+) samples. Total tau (Tau5) did not change between all groups. Next, we measured PAD2 and PAD4 expression by immunohistochemistry (IHC). IHC staining showed increased PAD2 immunoreactivity in astrocytes and increased neuronal labeling of PAD4 immunoreactivity in cortical AD tissue compared to healthy controls (**Figure 1C**). PAD2 and PAD4 expression was significantly increased in AT8 (+) human tissue as compared to the control samples and AT8 low (−) samples (**Figure 1D, E**). These data confirm increased PAD2 and PAD4 in AD brains similar to other reports but also indicates an increased association with pathological tau accumulation in a subset of patients.

**Figure 1.**
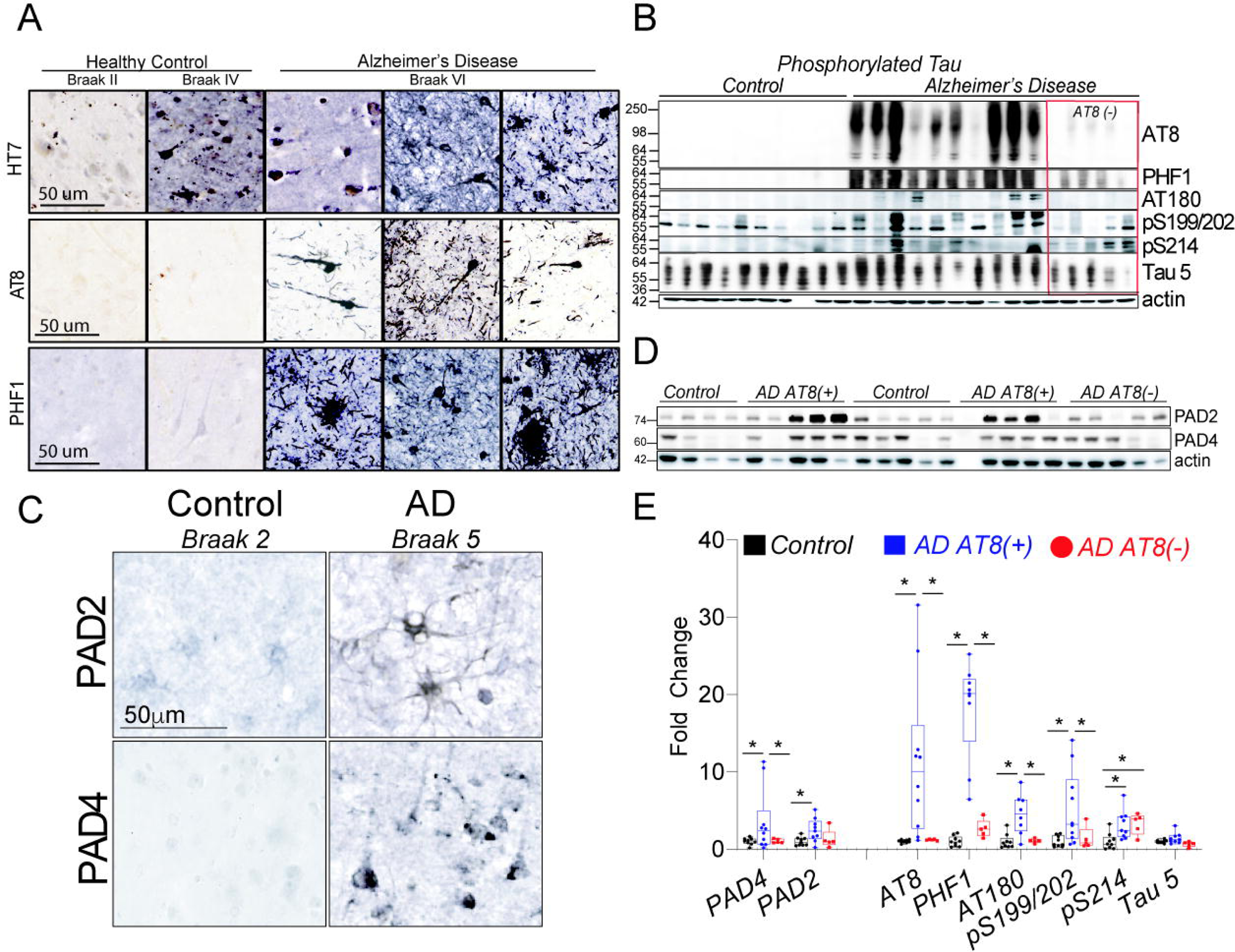
Tau and PAD expression in Alzheimer’s disease (AD) brain. **A**. Immunohistochemical images from healthy control samples Braak II-IV and AD samples Braak VI stained for total tau (HT7), AT8, and PHF1. **B**. Western blot analysis of AT8, PHF1 AT180, pS199/202, pS214, and total tau (Tau5) expression in cortex homogenates from AD brain and unaffected controls. AD samples were segregated into AT8 high (+) or AT8 low (−) groups. **C**. Shows western blot analysis and densitometry for tau epitopes from AD and healthy controls. **D**. Images show increased astrocytic PAD2 and neuronal PAD4 immunoreactivity in AD brain compared to the control tissue. **E**. Western blot analysis of PAD2 and PAD4 expression in hippocampal homogenates from AD brain and unaffected controls. AD samples were designated as either AT8 high (+) or AT8 low (−). PAD2 and PAD4 expression increased in AT8 high (+) samples compared to unaffected controls. Values were normalized to actin and graphed as fold change (± SEM). Statistical analysis performed by One-way Anova with Fisher’s LSD as post-hoc analysis (*P < 0.05, n=10/ control sample, n=15/ AD sample). Scale bar represents 50 μm

**Table 1.** Age and PMI mean ± standard deviation (SD) of AD and unaffected healthy control cases. Table 1 list de-identified samples and randomly re-assigned a sample (Case ID) based on the sample ID NIH Neurobiobank listed in for their tissue. Listed in the table also includes control vs. Alzheimer’s disease (AD), brain repository, clinical diagnosis, brain region, sex, year at death, and available Braak stage.

| Case ID | Brain Bank | Neuropathological Diagnosis | Tissue | Sex | Age at Death (Years) | Braak Stage |
| --- | --- | --- | --- | --- | --- | --- |
| NIH1-1 | Mt. Sinai | Control | Cerebral Cortex BM7 | F | 89 | 1 |
| NIH1-2 | Mt. Sinai | Control | Cerebral Cortex BM7 | F | 86 | 1 |
| NIH1-3 | Mt. Sinai | Control | Cerebral Cortex | M | 66 | 1 |
| NIH1-4 | Mt. Sinai | Control | Cerebral Cortex BM7 | F | 70 | 2 |
| NIH1-5 | Mt. Sinai | Control | Cerebral Cortex BM7 | M | 83 | 3 |
| NIH1-6 | Univ. Maryland | Control | Cerebral Cortex | M | 82 | N/A |
| NIH1-7 | Univ. Maryland | Control | Cerebral Cortex | F | 80 | N/A |
| NIH1-8 | Univ. Maryland | Control | Cerebral Cortex | M | 78 | N/A |
| NIH1-9 | Univ. Maryland | Control | Cerebral Cortex | F | 82 | N/A |
| NIH1-10 | Univ. Maryland | Control | Cerebral Cortex | M | 69 | N/A |
| NIH1-11 | Mt. Sinai | AD | Cerebral Cortex BM10 | F | 100 | 6 |
| NIH1-12 | Mt. Sinai | AD | Cerebral Cortex BM8 | F | 83 | 6 |
| NIH1-13 | Mt. Sinai | AD | Cerebral Cortex BM7 | M | 80 | 6 |
| NIH1-14 | Mt. Sinai | AD | Cerebral Cortex BM9 | F | 82 | 6 |
| NIH1-15 | Mt. Sinai | AD | Cerebral Cortex BM11 | M | 89 | 6 |
| NIH1-16 | Univ, Miami | AD | Primary Motor Cortex | M | 80 | 6 |
| NIH1-17 | Univ, Miami | AD | Primary Motor Cortex | F | 80 | 6 |
| NIH1-18 | Univ, Miami | AD | Primary Motor Cortex | F | 81 | 2-3 |
| NIH1-19 | Univ, Miami | AD | Primary Motor Cortex | M | 82 | 6 |
| NIH1-20 | Univ, Miami | AD | Primary Motor Cortex | M | 83 | 3-4 |
| NIH1-21 | Univ, Miami | AD | Primary Motor Cortex | M | 88 | 6 |
| NIH1-22 | Univ, Miami | AD | Primary Motor Cortex | M | 78 | 3 |
| NIH1-23 | Univ, Miami | AD | Primary Motor Cortex | M | 78 | 6 |
| NIH1-24 | Univ, Miami | AD | Primary Motor Cortex | F | 77 | 6 |
| NIH1-25 | Univ, Miami | AD | Primary Motor Cortex | F | 67 | 6 |

### Site-specific identification of Tau citrullination by mass spectrometry

Citrullination of arginine residues results in the addition of 0.984 Daltons, which can be resolved from unmodified arginine residues using high resolution and high accuracy mass spectrometry. We used total histone 3, which is a client of PAD4 and PAD2 to verify the citrullination reaction (**Figure 2A**). Using this approach, we performed the same reaction with 2N4R (441aa) full-length tau and subjected the reaction mixture to SDS-PAGE gel following incubation with a citrulline incorporating rhodamine-PG (Rh-PG) fluorescent probe (**Figure 2A**). We found increased fluorescent bands in the PAD2+tau and PAD4+tau reaction suggesting successful arginine to citrulline conversion within tau. Next, we subjected digested bands to mass spectrometry analysis to identify site specific citrullination of recombinant 2N4R (**Figure 2B**). Tau was citrullinated at all 14 arginine residues between either PAD4 or PAD2. To ensure the correct identification of the site-specific citrullinated tau, we verified six citR tau epitopes for sequence coverage for each citrulline modification containing the peptide within the MS/MS and y-ion series (i.e., tau citR 194 **Figure 3A, Supplemental Figure 1**). This confirmed the modified vs. unmodified epitope spectra for citrullination within tau protein (**Figure 2B**).

**Figure 2.**
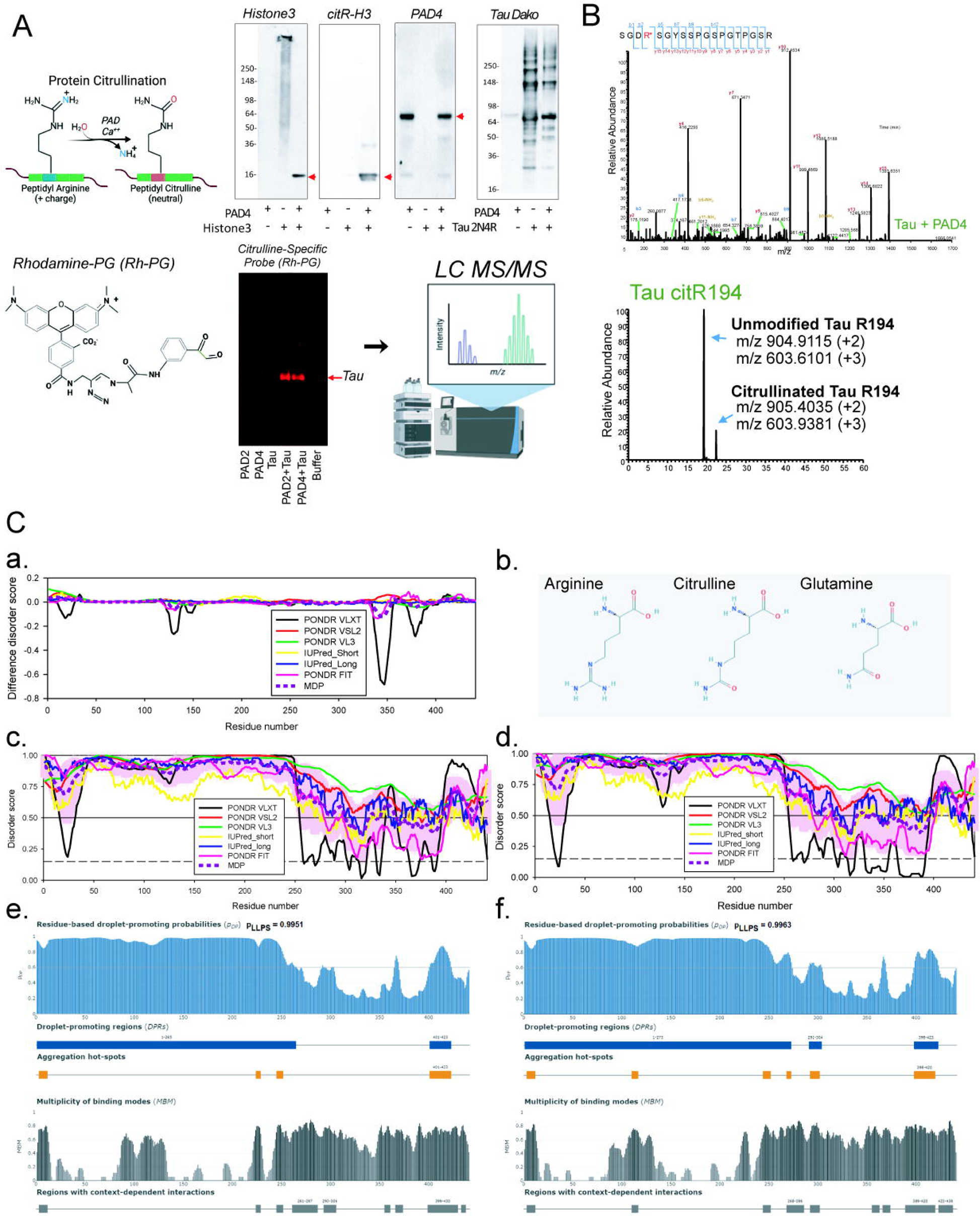
Identification of tau citrullination by mass spectrometry. **A**. Two micrograms of recombinant PAD4 with 1μg of recombinant 2N4R tau protein (441aa) or 1 μg histone 3 was incubated for 90 minutes. Western blot analysis showed citrullination of histone 3. Rhodamine - phenyl-gloxal (Rh-PG) fluorescent indicator probe was used to detect citrulline incorporation into protein. Rh-PG showed the PAD2 and PAD4 incubation with tau increased fluorescent incorporation. **B**. Samples were gel extracted and subjected to mass spectrometry to identify tau citrullination. In total, 14 arginine sites were citrullinated on full length tau 2N4R (441aa). The y-ion series of tau citR194 was detected in the region of the modification and confirmed by the modified vs. unmodified citR194 epitope spectra. **C** Analysis of the effect of the pseudo-citrullination on intrinsic disorder0related properties of tau. **C**.**a**. “Difference disorder spectrum” generated by the subtraction of the per-residue disorder propensities generated by the individual predictors for the normal tau are subtracted from the corresponding profiles generated for its pseudo-citrullinated R→Q form. **b**. Chemical structures of arginine, citrulline and glutamine. **c**. Multifactorial per-residue disorder profile generated for the normal tau using a set of commonly used disorder predictors. **d**. Multifactorial per-residue disorder profile generated for the pseudo-citrullinated R→Q tau using a set of commonly used disorder predictors. Plots **C.c**. and **C.d**. were generated using the RIDAO web platform that combines the outputs of PONDR^®^ FIT, PONDR^®^ VSL2, PONDR^®^ VL3, PONDR^®^ VLXT, IUPred Short, and IUPred Long to generate an integral disorder profile of a query protein. For each protein, RIDAO also calculates the mean disorder profile (MDP) by averaging the outputs of all individual predictors. The disorder score was assigned to each residue, with a residue with disorder score equal to or above 0.5 being considered as disordered and a residue with disorder score below 0.5 being predicted as ordered. Residues/regions with disorder scores between 0.15 and 0.5 were considered as ordered but flexible. **e**. The FuzDrop-generated plot generated for normal tau and showing the sequence distribution of the residue-based droplet-promoting probabilities, p_DP_ (upper panel) and the multiplicity of binding modes plot showing positions of regions that can sample multiple binding modes in the cellular context (sub-cellular localization, partners, posttranslational modifications)-dependent manner (bottom panel). **f**. LLPS potential and MBM plots generated for the pseudo-citrullinated R→Q tau by FuzDrop.

**Figure 3.**
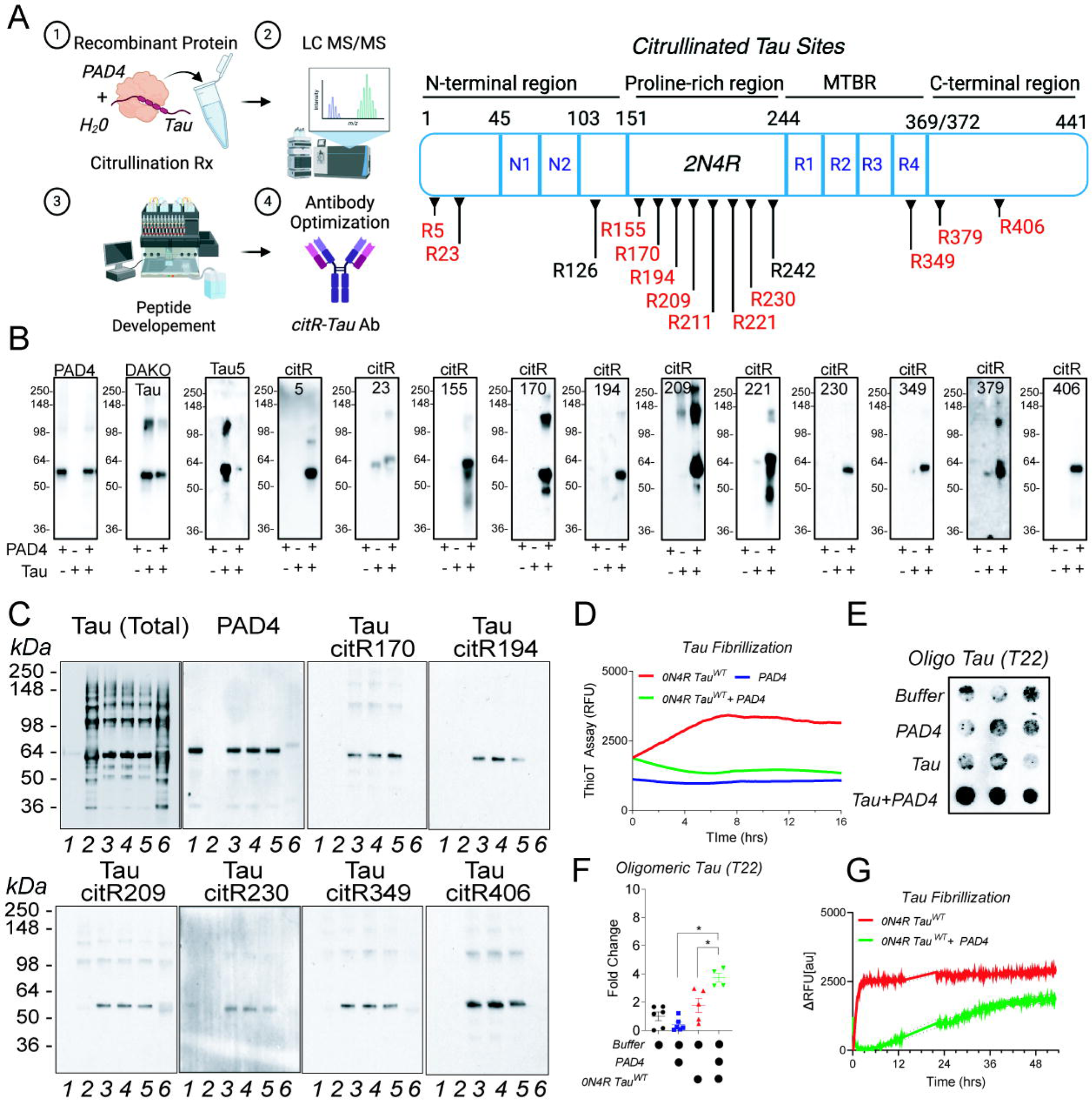
Development of novel citrullinated tau antibodies. **A.** Schematic of citR epitope positions within the tau (441a.a.) protein sequence. Eleven novel antibodies were generated against citR tau. **B.** Western blot shows *in vitro* recognition of PAD4, total tau (DAKO tau, rabbit polyclonal), total tau (Tau5, monoclonal antibody epitope sequence 210-241), tau citR5, tau citR23, tau citR155, tau citR170, tau R194, tau R209, tau citR221, tau R230, tau R349, tau citR379 and tau citR406 tau following enzymatic reaction of recombinant tau and PAD4. **C**. *In vitro* citR tau abolished by pharmacological inhibition of PAD4. Enzymatic reaction between recombinant PAD4 and 4R2N tau in 1 : 1 molar ratio with increasing concentrations of Ca^2^ [2-10mM] or 300 μM BB-Cl-amidine. Western blot analysis of total tau, PAD4 and tau citR170, tau cit194, tau citR209, tau citR230, tau citR349, tau citR406 demonstrated the citR tau antibodies specificity to citR tau; PAD4 (lanes 1, 3-6), tau (lanes 2-6), Ca^2+^ (lanes 3-6) induced citR-tau (lane 3-5). PAD4 inhibitor (BB-Cl-amidine) (lane 6) inhibited all citR-tau epitopes. Note that total tau levels did not change following BB-Cl-amidine treatment. **D**. Thioflavin T (ThT) assay of tau fibrillization following incubation with PAD4 (2 μg, blue line) in the presence of 1: 4 ratio with heparin (8 μg). Tau fibrillization increased over 16 hours however citrullinated tau failed to aggregate after 16 hours revealed that citrullination halts tau fibrillization. PAD4 alone (blue line) did not aggregate. **E-F.** Aliquots were analyzed by dot blot and probed with oligomeric tau specific T22 antibody. Densitometry analysis of dot intensity showed significant increased tau oligomerization of citrullinated tau. **G**. Tau fibrillization extended over 48 hours showed that after 20 hours citrullinated tau began to aggregate to a lesser extent compared to tau alone.

Little is known about the functional state of citrullinated tau. Given the charge neutralization, increased hydrophobicity, and reduced π-π interactions, we sought to understand the effect of citrullination on intrinsic disordered status of tau using bioinformatics tools. There are no natural codons that code for citrulline, and this residue, being not common and well-characterized in protein structures, is not included in any computational analytic platforms. Therefore, to mimic the potential citrullination effects, we changed all arginine (R) residues to glutamine (Q) residues in the 2N4R tau as pseudomimetics for citrulline and used a set of computational tools designed for the intrinsic disorder analysis. This all R-to-Q substitution in tau caused a slight increase in the overall mean hydrophobicity (from 0.4036 to 0.4071 in Kate-Doolittle hydropathy scale (Kyte and Doolittle 1982) of the protein. At the same time, the total protein charge changed from +2 to −12, leading to the increase in the absolute mean net charge from 0.0045 to 0.0272. As a result, position of a pseudo-citrullinated R→Q tau within the charge-hydropathy (CH) plot (Uversky, Gillespie et al. 2000, Oldfield, Cheng et al. 2005) shifted closer to the boundary separating disordered and ordered proteins (distance to the boundary decreased from 0.0273 to 0.0160, data not shown), indicating that citrullination might lead to a slight decrease in the overall disorder predisposition of tau. This is an interesting, conflicting, and not an obvious conclusion, since pseudo-citrullination shows opposite effects on tau hydrophobicity and mean net charge, according to the logic behind the CH-plot analysis, increased hydrophobicity should lead to the decrease in disorder propensity, whereas increased mean net charge is typically associated with the increase in disorder potential. This conclusion is further supported by the evaluation of the per-residue disorder propensity to tau and pseudo-citrullinated R→Q tau by a set of commonly used disorder predictors assembled using the RIDAO platform (Dayhoff and Uversky 2022). Results of these analyses are summarized in **Figure 2C** representing the multifactorial disorder profiles for the normal and pseudo-citrullinated R→Q tau and also showing the “disorder difference spectrum”, where the disorder profiles generated by individual predictors for the normal tau are subtracted from the corresponding profiles generated for its pseudo-citrullinated form. The mean disorder characteristics (percent of predicted intrinsically disordered residues (PPIRD) and average disorder score (ADS)) generated by these tools provide further evidence of the “confusing” outputs of pseudo-citrullination. In fact, the normal tau is characterized by PPIDR-ADS of 65.99%-0.6714 (PONDR^®^ VLXT), 98.41%-0.8161 (PONDR^®^ VSL2), 100.00%-0.8462 (PONDR^®^ VL3), 76.42%-0.6486 (IUPred_Short), 90.02%-0.7848 (IUPred-Long), 72.11%-0.7110 (PONDR^®^ FIT), and 83.90%-0.7463 (MDP). At the same time, these tools generated the following outputs for the pseudo-citrullinated R→Q tau: 63.27%-0.6322 (PONDR^®^ VLXT), 99.32%-0.8306 (PONDR^®^ VSL2), 100.00%-0.8526 (PONDR^®^ VL3), 76.42%-0.6575 (IUPred_Short), 88.21%-0.7864 (IUPred-Long), 72.34%-0.7107 (PONDR^®^ FIT), and 81.63%-0.7450 (MDP). It is also important to emphasize here that despite these subtle differences, the protein is still predicted as mostly disordered irrespectively of its pseudo-citrullination status.

Since tau is known to undergo liquid-liquid phase separation (LLPS), at the next step, we looked on how pseudo-citrullination might affect the LLPS potential of this protein. Results of the corresponding analysis by FuzDrop (Hardenberg, Horvath et al. 2020, Vendruscolo and Fuxreiter 2022) are shown in **Figure 2C**. This analysis provided several interesting observations. First, both forms of proteins are capable of spontaneous LLPS, with pseudo-citrullinated tau being a bit more potent droplet driver (its probability of spontaneous LLPS is p_LLPS_ = 0.9963 versus 0.9951). Normal tau has two droplet-promoting regions (DPRs, residues 1-265 and 401-423), as well as four aggregation hot spots (residues 3-12, 224-229, 245-252, and 401-423); i.e., regions that might drive protein aggregation within the droplets. There are also 9 regions with context-dependent interactions or regions with multiplicity of binding modes (MBM, residues 3-12, 224-229, 245-252, 261-287, 293-306, 355-361, 366-374, 399-430, and 433-438); i.e., regions that exhibit different binding modes depending on the peculiarities of cellular context (sub-cellular localization, partners, posttranslational modifications, etc.). On the other hand, there are three DPRs (residues 1-273, 291-304, and 398-423) in the pseudo-citrullinated R→Q tau, which also contains 6 aggregation hot spots (residues 3-12, 110-117, 244-252, 268-273, 292-302, and 398-420) and 9 MBM regions (residues 3-12, 110-117, 244-252, 268-286, 292-302, 355-363, 366-374, 389-420, and 423-438). These results indicate that citrullination might affect LLPS potential of tau and also can enhance the aggregation predisposition of tau, since it results in doubling of the number of aggregation hot spots.

Based on these outputs of the computational analyses, one might suggest that pseudo-citrullination increases hydrophobicity of tau and renders this protein slightly more ordered and more prone for spontaneous LLPS. It also furnishes the protein with higher aggregation potential likely due to the increased potential to form oligomers due to the increased local hydrophobicity.

### Development of a citrullinated tau antibody panel

To confirm multi-epitope citrullination of tau we developed a panel of affinity purified citR tau antibodies to the N-terminal domain (tau citR5, citR23), the proline rich region (tau citR155, citR170, citR194, citR209, citR221, citR230), the microtubule binding repeat domain (tau citR349) and the c-terminal domain (tau citR379, citR406) (**Figure 3A-B**). We validated antibodies with recombinant tau 441 and PAD4 enzymatic reactions (as previously described). While untreated recombinant tau or PAD4 alone was not detected by any of the eleven antibodies, strong labeling was observed for all 11-modified citrullinated tau epitopes (**Figure 3B**). PAD4, total tau (DAKO tau, rabbit polyclonal), and Tau5 (clone, mouse) were used to confirm expression levels of PAD4 and 2N4R tau. While DAKO tau antibody labeled total tau in both lanes (unmodified and modified citR tau), Tau5 failed to label the modified citR tau reaction (**Figure 3B**). This is noteworthy because Tau5 binds the tau epitope sequence 210-241, which harbors three citrullinated tau epitopes within this sequence (i.e., tau citR 211, tau citR 221, tau citR230) and two epitopes flanking the Tau5 sequence (i.e., tau citR 209 and tau citR242). These data suggest that citrullination of tau is sufficient to disrupt Tau5 antibody binding. Given the potential extent of tau citrullination, it is worth considering that certain tau immunotherapies may show reduced efficacy to tau because of post-translational modifications like citrullination that could reduce immunotherapy interactions.

Next, we validated the specificity of the enzyme reaction by testing 6 citR tau antibodies (citR179, citR194, citR209, citR230, citR349, citR406) against pharmacological inhibition of PAD4. Citrullination reactions between PAD4 and 2N4R tau were performed with increasing concentrations of Ca^2^ [2-10mM] or 300 μM BB-Cl-amidine (irreversible PAD4 inhibitor). Western blot analysis confirmed that all 6 citR tau antibodies recognized citR-tau (lane 3-5) but not PAD4 (lanes 1) or tau alone (unmodified) (lanes 2) (**Figure 3C**). However, in the presence of BB-Cl-amidine (lane 6) citrullination was inhibited at all the citR-tau epitopes tested (**Figure 3C**). Notably, increased [Ca^2+^] did not affect citR tau levels (lanes 3-6), suggesting that 2mM Ca^2+^ was sufficient to maximize PAD4-mediated citrullination of tau citR. These data suggest that citrullination occurs throughout multiple domains of tau under recombinant protein systems, and that our panel of antibodies detected PAD4-mediated tau citrullination. Finally, the citR tau modification can be blocked by PAD inhibitors.

We also tested whether PAD4-induced citrullination impacted tau aggregation through fibrillization via beta sheet formation using Thioflavin T assay. Tau (0N4R) was citrullinated with PAD4 for 90 minutes prior to fibrillization monitoring for 16 hours. Tau indeed began to form fibrils through ThioT incorporation within 4 hours following aggregation conditions, however citrullinated tau failed to produce any ThioT fluorescence or fibrils under these conditions (**Figure 3D**). PAD4 alone under the same conditions also failed to aggregate suggesting low or no intrinsic self-assembly toward forming an amyloid. After 16 hours we tested conditioned samples for signs of tau oligomers using anti-oligomeric tau T22 antibody. Interestingly, citrullinated tau showed significantly more oligomers than tau alone, which were converted to fibrils (**Figure 3E, F**). Next, we extended the time of aggregation under these conditions in follow up experiments. Similarly, tau alone began to form ThioT fibril aggregation for 3 hours, however citrullinated tau aggregation was significantly delayed but began to show ThioT positive fibrils after 16 hours (**Figure 3G**). These data coupled with the interpretation of pseudo-citrullinated tau from computer simulations suggest that citrullination of tau slows fibrillization rate at the expense of accumulating citrullinated tau oligomers.

### Calcium load and PAD4 expression regulate substrate specificity for citR tau epitopes

An important yet untested hypothesis for citrullination via PAD activity, and perhaps the most biologically intriguing mechanism, is that PAD4 displays different “*coordinated activation states*” depending on its five Ca^2+^ binding pockets. Low Ca^2+^ load may display high substrate specificity (i.e., restricted epitope citrullination), whereas high Ca^2+^ binding capacity displays low substrate specificity (i.e., non-restricted citrullination). An alternative to non-restricted citrullination may also depend on PAD concentration itself. If true, this would suggest that certain regions within tau may undergo citrullination at different rates depending on calcium loads, PAD concentration and pathological stages (aberrant Ca^2+^ dysfunction) of disease. Given our panel of citrullinated tau antibodies and the ability to measure tau at multiple functional domains, we sought to test this by monitoring the degree of tau citrullination at six citR tau sites (i.e., tau citR170, citR 194, citR 209, citR230, citR349, citR406) under various PAD4 concentrations (0.67-670nM) with high calcium load (10mM) or without calcium (**Figure 4A-B**). Indeed, PAD4 concentration (670nM) with high calcium load (10mM) citrullinated all six tau sites tested. Interestingly PAD4 (670nM) with no calcium also citrullinated tau at citR194, citR209, citR230, and citR349 equally, however citrullination of citR170 and tau citR406 was diminished when calcium was absent suggesting that certain citR epitopes are sensitive to coordinated calcium activation of PAD4. Importantly, a 10-fold lower concentration of PAD4 (67nM) with high calcium load (10mM) was also able to restore citrullination of tau citR170 and tau citR406 beyond that of PAD4 (670nM +0mM calcium) also indicating the degree of PAD4 specificity for selective epitopes. As PAD4 concentration decreased, citrullination of tau also decreased at all epitopes. These data suggest that citrullination of tau depends on both PAD4 concentration and calcium load but can vary in epitope specificity depending coordinated calcium activation of PAD4. This is a notable finding for an irreversible modification because it dictates how a limited number of PADs expressed in the brain regulate many proteins with substrate specificity.

**Figure 4.**
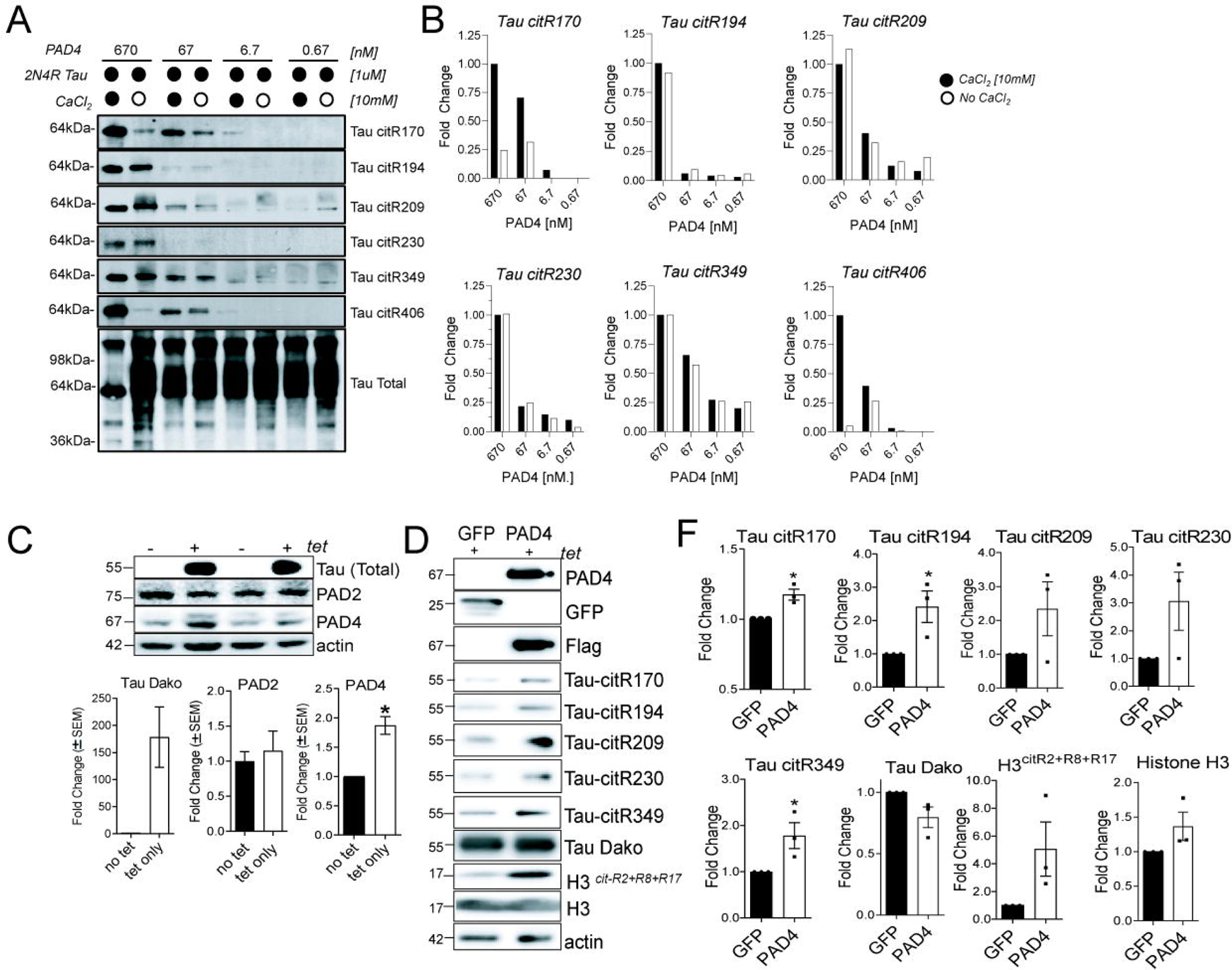
Tau citrullinated epitope specificity is dependent on PAD4 concentration and calcium concentration. **A**. Enzyme reaction of citrullinated tau was performed using 1uM tau, various concentrations of PAD4 (0.67-670nM) with/ or without 10mM CaCl_2_. And probed for tau citR170, tau citR194, tau citR209, tau citR349, tau citR406, and total tau. **B**. showed densitometry at each PAD4 concentration. Data shows that tau citR170, and tau citR406 were calcium dependent, whereas tau citR194, tau citR209, tau citR230, tau citR 349 were calcium independent but citrullinated at higher PAD4 concentrations. **C**. Tau induces PAD4 expression in iHEK stable tau cells. Western blot analysis showed that tau induction upon tetracycline (tet) treatment increased PAD4 expression but not PAD2 expression. **D** iHEK tau cells transfected with GFP or PAD4-flag plasmid DNA for 72hrs showed increased tau citR170, tau citR194, tau citR209, tau citR230, and tau citR349 levels during PAD4 overexpression compared to GFP expressing cells. PAD4 overexpression also increased citrullinated histone 3 (citR-H3 R2+R8+R17). **F**. Represents densitometry values normalized to actin and graphed as fold change (± SEM). Statistical analysis performed by Student’s t-test (*P < 0.05, n=3).

To understand the bidirectional relationship between tau induction and PAD expression we utilized the inducible tau cells iHEK-tau cell line as previously described (Abisambra, Jinwal et al. 2013). The administration of tetracycline in the iHEK-tau cells media induced total tau expression. Quantitative analysis showed that tet-induced tau expression significantly increased PAD4 but not PAD2 expression in these cells (**Figure 4C**). We also tested if PAD4 overexpression impacted citR tau in iHEK-tau cells. Tet-induced iHEK tau cells were transfected with GFP or PAD4-flag plasmid DNA for 72hrs. Indeed, PAD4 overexpression increased tau citR170, citR194, citR209, citR230, and citR349 (others citR tau not tested) levels compared to GFP expressing cells (**Figure 4D-F**). However, it should also be noted that citR tau was detected at low levels in GFP expressing cells. PAD4 activity was further confirmed by increased levels of citrullinated histone 3 (citR-H3 R2+R8+R17) compared to GFP transfected cells (**Figure 4D-F**). These data suggest a reciprocal relationship between tau biology and PAD4 activity but also that tau citrullination occurs in cellular models.

### PAD2 and PAD4 expression is increased in mouse models of tauopathy

To confirm CNS expression of PAD enzymes, we stained rTg4510, PS19 mice and non-transgenic littermates for PAD4. PAD4 increased in the hippocampus of both tauopathy models (**Figure 5A**) along with citrullinated histone 3 (citR-H3) (**Figure 5B, C**), a substrate for PAD4 activity (Wang, Li et al. 2009). PAD2 increased in glia (astrocytes), while PAD4 increased in neurons and glia (**Figure 5D**). Next, we measured the temporal profile of both PAD2 and PAD4 in young and aged rTg4510 mice and non-transgenic littermates. Cortical homogenates from 4-, 8-, 13- and 16-month-old rTg4510 mice and age-matched non-transgenic littermates were measured for PAD2 and PAD4 expression levels by Western blot (**Figure 5E-G**). Densitometry analysis showed increased in PAD2 levels in aged 13- and 16-month-old rTg4510 mice compared to young rTg4510 mice and non-transgenic littermates (**Figure 5E, F**). Conversely, PAD4 levels were significantly increased in young 4-month-old rTg4510 mice compared to age-matched non-transgenic littermates (**Figure 5E, G**). These data suggest early expression of PAD4 during tau pathogenesis, whereas PAD2 increased at later stages, possibly suggesting glial reactivity in response to the advanced tau phenotype.

**Figure 5.**
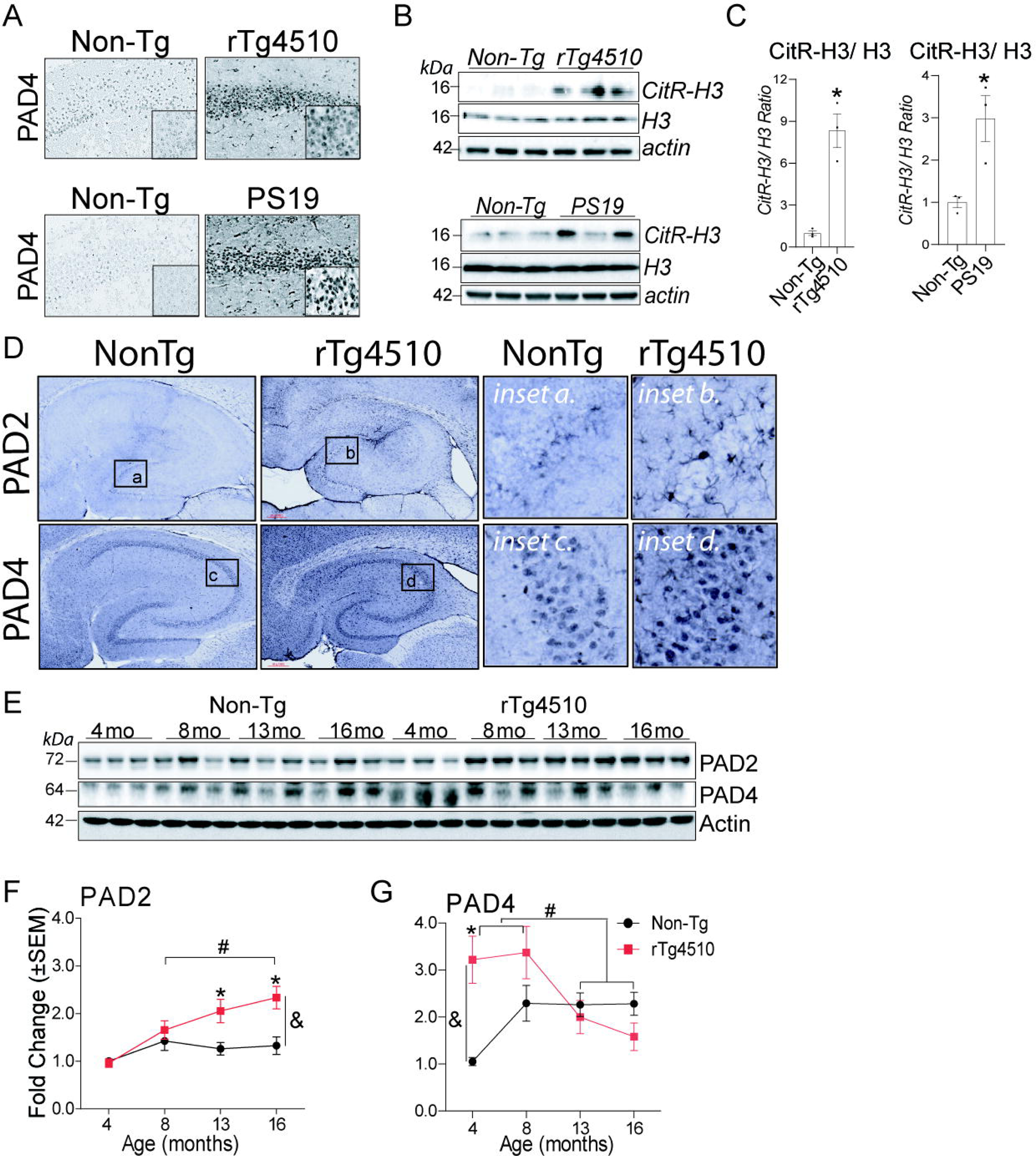
PAD expression and activity is increased in mouse models of tauopathy. **A.** Immunohistochemical images of rTg4510, PS19 tau mice and non-transgenic (NonTg) littermates in hippocampi tissue labeled against PAD4 showed increased neuronal PAD4 immunoreactivity in tau mice brain compared to the NonTg tissue. **B** Citrullination activity was measured biochemically in the brain homogenates probed for citR-H3 and total H3 antibodies. **C**. Citrullinated histone3 levels (citR-H3) increased in rTg4510 and PS19 tissue compared to age-matched Non-Tg littermates. Values were normalized to histone 3 (H3) and graphed as fold change (± SEM). Statistical analysis performed by student t-test (*P < 0.05, n=3). **D**. PAD2 and PAD4 expression in tauopathy (rTg4510 tau transgenic) mice and non-transgenic littermates show astrocytic PAD2 staining and neuronal and glial PAD4 staining in the hippocampus. **E**. Panel shows PAD2 and PAD4 expression in tauopathy mice aged for 4, 8, 13, and 16 months. Statistical analysis were performed by two-way ANOVA; &=denotes significance by genotype, # =by age, * =pairwise comparison between genotype within the age group, ^+^ =denotes pairwise comparison of rTg4510 across age P < 0.05 (n=3-8/ age).

### Induction of citrullinated tau in animal models of tauopathy

To gain further insights on the degree to which citR tau occurs in animal models of tauopathy, we performed immunohistochemical (IHC) and western blotting utilizing the 11 antibody panel if citR tau including (tau citR5, tau citR23, tau citR155, tau citR170, tau citR194, tau citR209, tau citR221, tau citR230, tau citR349, tau citR379, tau citR406) in three tau models; rTg4510 (*MAPT P301L, 0N4R*), PS19 (*MAPT P301S, 1N4R*) and 4RTg2652 mice (*MAPT wild-type tau, 1N4R*) transgenic mouse models (Wheeler, McMillan et al. 2015)). Hippocampal images of 8-month-old rTg4510 tau mice confirmed increased immunoreactivity for all eleven citR tau epitopes with the exception the tau citR406 epitope compared to age-matched non-transgenic littermates (**Figure 6A**). The high magnification of CA3 region also demonstrated neuronal expression for citR tau except citR406 in neurons compared to non-transgenic littermates (**Figure 6A**). We also validated citR tau antibodies via western blot and confirmed increased expression of all citR tau epitopes again with the exception of tau citR406 in 8-month-old rTg4510 mice, PS19 and 4RTg2652 mice compared to their respective non-transgenic littermates (**Figure 6B**). These data signify increased tau citrullination in multiple animal models of tauopathy but also indicates that citR tau is a common PTM that occurs during tau pathogenesis. Further work is necessary to understand why tau citR406 does not show the same profile as other citR tau epitopes in all three tauopathy models but can become citrullinated with recombinant protein systems.

**Figure 6.**
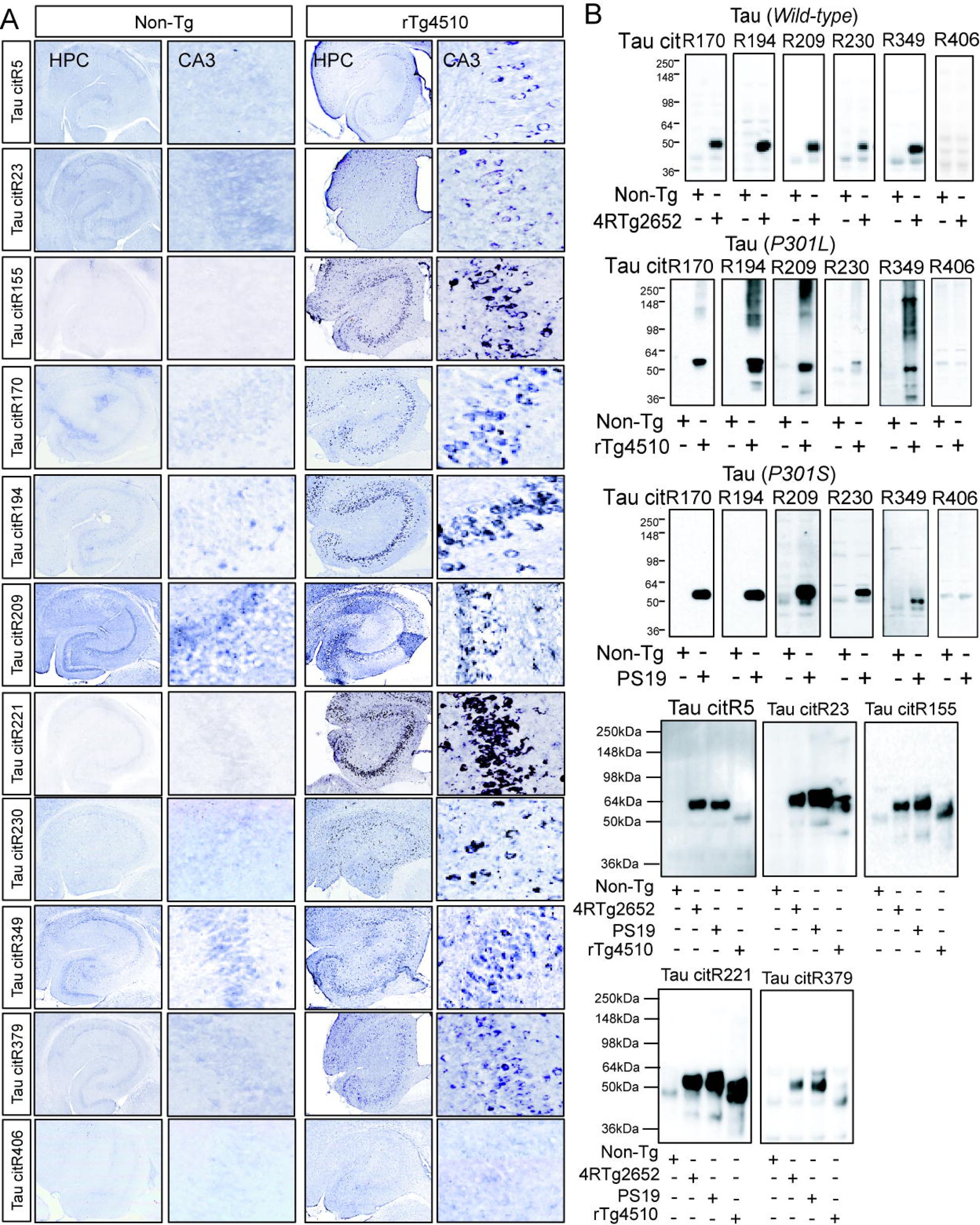
Citrullinated tau is increased in multiple animal models of tauopathy. Immunohistochemical images of rTg4510 and non-transgenic (NonTg) littermates in hippocampal tissue labeled against citrullinated tau. **A.** Shows increased neuronal expression of tau citR5, tau citR23, tau citR155, tau citR170, tau citR194, tau citR209, tau citR221, tau citR230, tau citR349, tau citR379, and little to no expression of tau citR406 in rTg4510 mice compared to the Non-Tg littermate tissue. Panels show hippocampus (HPC) and CA3 region (magnified panel). **B.** Shows western blot images of increased tau citR5, tau citR23, tau citR155, tau citR170, tau citR194, tau citR209, tau citR221, tau citR230, tau citR349, tau citR379, and little to no expression of tau citR406 in rTg4510 (*MAPT P301L; 0N4R*), PS19 (*MAPT P301S; 1N4R*), and 4RTg2652 (*MAPT wildtype; 1N4R*) mice compared to Non-Tg littermates.

### Aging analysis of tangle pathology and citrullinated tau in tauopathy mice

To determine the temporal expression of citR tau, we performed immunohistochemistry in rTg4510 mice at 4-, 8-, 13-, 16-months old mice and non-transgenic littermates (**Figure 7A**). Tissue from each age group was stained for human tau (HT7 antibody) to determine the regional tau burden. Images of anterior cortex (ACX), hippocampus (HPC) and entorhinal cortex (ECX) and magnified insets showed total tau deposition at all ages (**Figure 7B**). The quantification of percent area of positive staining showed increased total tau levels at 4 months, which peak at 8-months and waned at 13- and 16-month in each region (**Figure 7B**). Decreased tau at 13- and 16-month-old possibly reflects ongoing neurodegeneration in this model. Non-transgenic mice lacked immunoreactivity for total tau (HT7) at any age. Next, we performed Gallyas Silver staining on non-transgenic and rTg4510 mice to measure neurofibrillary tangle (NFT) pathology (**Figure 7C**). The quantification of percent area showed no tangle pathology at 4-months, progressively increased at 8-months and peaked at 13-months, while slight decreasing at 16-months in the ACX, HPC and ECX of rTg4510 mice (**Figure 7C**). Non-transgenic mice displayed no reactivity to Gallyas silver staining at any age. These data confirm age-related tangle accumulation in rTg4510 mice.

**Figure 7.**
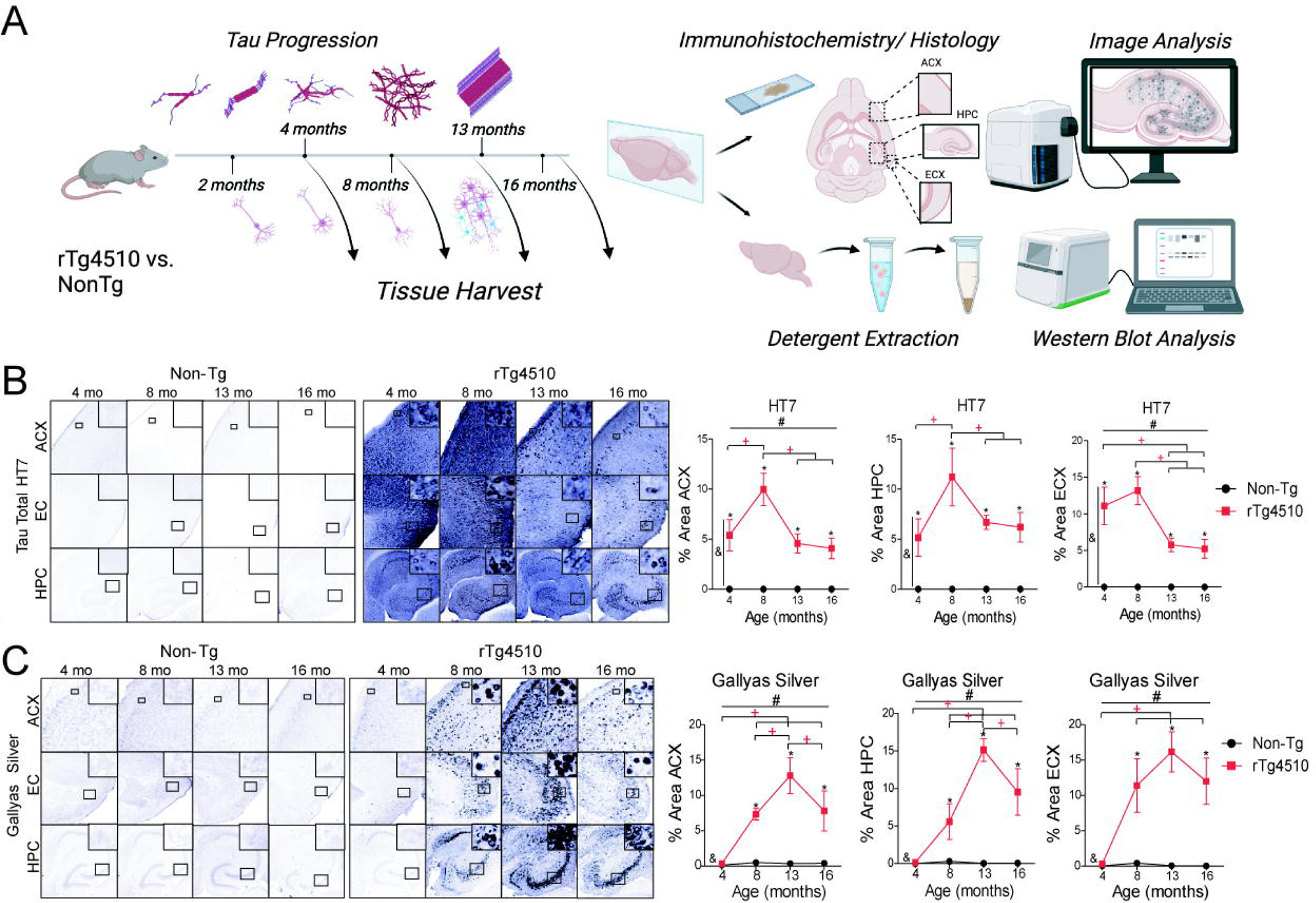
Aging analysis of tau neurofibrillary tangle progression in rTg4510 mice. **A**. Schematic of tau progression and neurodegeneration in rTg4510 mice. **B.** Tissue from non-transgenic mice and rTg4510 mice at 4, 8, 13, and 16 months immunohistochemically stained for total tau in the anterior cortex (ACX), hippocampus (HPC) and entorhinal cortex (ECX). **C**. Mice were histologically labeled for neurofibrillary tangles with Gallyas Silver stain in the ACX, HPC, ECX. Staining intensity analyzed as percent positive area showed increased total tau at all ages and tangle pathology developed by 8-13 months in ACX, HPC, and ECX. Statistical analysis were performed by two-way ANOVA; &=denotes significance by genotype, # =by age, * =pairwise comparison between genotype within the age group, ^+^ =denotes pairwise comparison of rTg4510 across age P < 0.05 (n=3-8/ age).

Next, we measured citR tau during pathological tau progression, by immunohistochemical analysis of citR tau antibodies (citR170, R194, R209, R230, and R349) in the anterior cortex (ACX), hippocampus (HPC), and entorhinal cortex (ECX). In general, 4-month-old rTg4510 mice showed higher levels of diffuse staining of citR170, R194, R209, R230, and R349 compared to non-transgenic littermates (**Figure 8**). Images from ACX, HPC and ECX regions demonstrated increased presence of immunoreactivity for citR170 (**Figure 8A-B**) and citR194 epitopes (**Figure 8C-D**) in neurons (magnified insets) with age. Image analysis showed significant expression peaking at 8-months of age for tau citR170 (**Figure 8A-B**) and tau citR194 (**Figure 8C-D**) in rTg4510 mice compared to non-transgenic littermates. Although expression of both tau citR170 and tau citR194 decreased compared to 8 months, their levels remained significantly elevated at 13- and 16-month-old rTg4510 mice compared to littermate controls. No citR tau immunoreactivity was detected in non-transgenic mice at any age.

**Figure 8.**
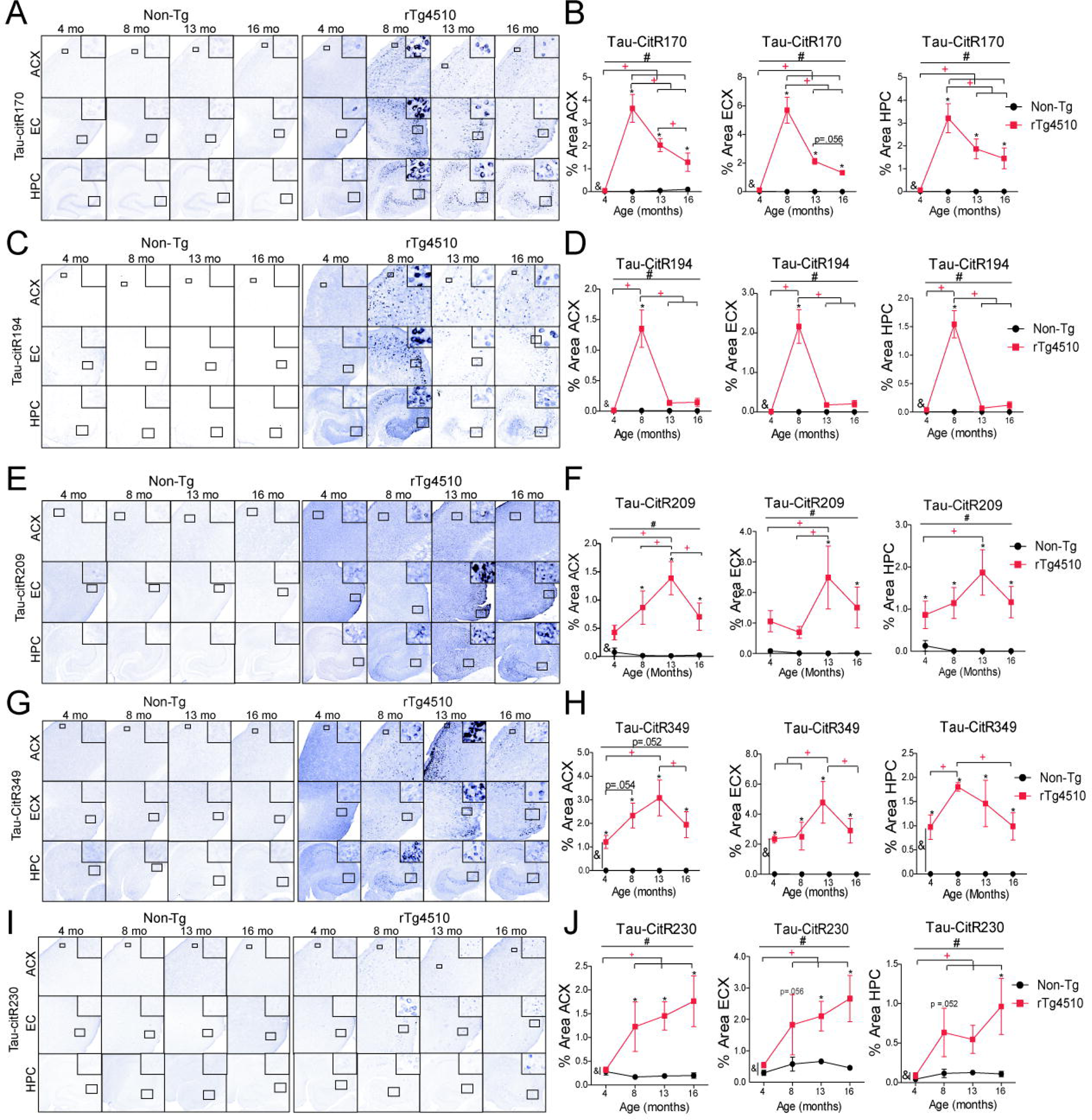
Citrullinated tau levels increase in animal models of tauopathy. **A** Immunohistochemistry images confirmed increased immunoreactivity in the anterior cortex (ACX), hippocampus (HPC) and entorhinal cortex (ECX) for tau citR170 (**A-B**), tau citR194 (**C-D**), tau citR209 (**E-F**), tau citR349 (**G-H)**, and au citR230 (**I-J**) in of rTg4510 tau mice compared to Non-Tg littermates at 4, 8, 13, and 16 months old (n=3-4/genotype). Age-related quantification of citrullinated tau and percent area for citR170 (**B**), tau citR194 (**D**), tau citR209 (**F**), tau citR349 (**H)**, and au citR230 (**J**) was performed. Statistical analysis were performed by two-way ANOVA; &=denotes significance by genotype, # =by age, * =pairwise comparison between genotype within the age group, ^+^ =denotes pairwise comparison of rTg4510 across age P < 0.05 (n=3-8/ age).

Similarly, we measured tau citR209 immunoreactivity (**Figure 8E-F**) in 4-, 8-, 13-, 16-month-old rTg4510 mice and non-transgenic littermates. Tissue from HPC of rTg4510 mice demonstrated increased neuronal immunoreactivity at all ages including 4-, 8-, 13-, and 16-months of age compared to non-transgenic mice (**Figure 8E-F**). The staining intensity (% positive area) showed significant tau citR209 immunoreactivity in ACX and ECX at 8- and 13-months, respectively peaking at 13-months (**Figure 8E-F**). Immunohistochemical analysis also demonstrated increased tau citR349 with age, starting at 4-months, peaking at 8-months in the hippocampus and 13-months in the anterior cortex and entorhinal cortex (**Figure 8G-H**). No citR labeling was detected in non-transgenic mice with age.

Finally, tau citR230 expression was comparable to non-transgenic controls at 4-months but increased immunoreactivity in neurons of aged rTg4510 tissue (**Figure 8I-J**). Magnified images (insets) demonstrated immunoreactivity of tau citR230 in neurons of each region. Interestingly, tau citR230 progressively accumulated with age. Although expression was low, positive staining in cortical regions (ACX, ECX) increased at 8-months and 13-months, respectively, and peaked at 16-month-old compared to non-transgenic littermates. Tau citR230 immunoreactivity failed to increase in the HPC until at 16-month-old in rTg4510 mice (**Figure 8I-J**). These findings suggest that the tau citrullination at various epitopes can occur at different stages when measured by immunohistochemistry.

### Biochemical analysis of phospho and citrullinated tau progression in tauopathy mice

We sought to understand to what degree and which biochemical fraction citrullinated tau accumulates during pathological progression. We performed western blot analysis on tau extracted via detergent free (TBS-soluble), sarkosyl soluble and sarkosyl insoluble methods in mice aged 4-, 8-, 13-, and 16-months in rTg4510 mice and non-transgenic littermates. We focused on tau at 55kDa, 64kDa and high molecular weight tau (98-148kDa) because these molecular species have indicated pathological tau transitions in this tauopathy model (Berger, Roder et al. 2007). Overall, we measured TBS-soluble total tau (DAKO), AT8, tau pSer214, tau citR170, citR194, citR209, citR230, and citR349 in mice. In general, all soluble tau epitopes measured showed a main effect of genotype and aged (**Figure 9**). Specifically, soluble total tau (**Figure 9A, B**), pSer214 (**Figure 9A, D**), tau citR170 (**Figure 9A, E**), citR194 (**Figure 9A, F**), citR209 (**Figure 9A, G**), citR230 (**Figure 9A, H**), and citR349 (**Figure 9A, I**) (55kDa), increased at 4-months of age but subsequently decreased by 8-16 months (with the exception tau citR209 at 16-months) compared to 4-month-old tau mice. Tau AT8 (although difficult to discriminate 55kDa from 64kDa) increased at 8-months of age and remained elevated throughout 16-months (**Figure 9A, C**). Total tau (**Figure 9A, B**), tau pSer214 (**Figure 9A, D**), and tau citR230 (**Figure 9A, H**) (64kDa) increased at 4-months and remained elevated throughout 16-months, whereas tau citR170 (**Figure 9A, E**), tau citR194 (**Figure 9A, F**), and tau citR349 (**Figure 9A, I**) increased at 8-months and remained elevated. Soluble total tau (98-148kDa) (**Figure 9A, B**), tau pSer214 (**Figure 9A, D**), and tau cit230 (**Figure 9A, H**) increased at 4-months and remained high, whereas tau citR170 (**Figure 9A, E**), tau citR194 (**Figure 9A, F**) increased by 4-months and waned by 16-months. Tau citR209 (98-148kDa) (**Figure 9A, G**) also increased by 4 months but decreased by 8-months. Finally, tau citR349 (98-148kDa) increased at 8-months and progressively accumulated through 16-months (**Figure 9A, I**).

**Figure 9.**
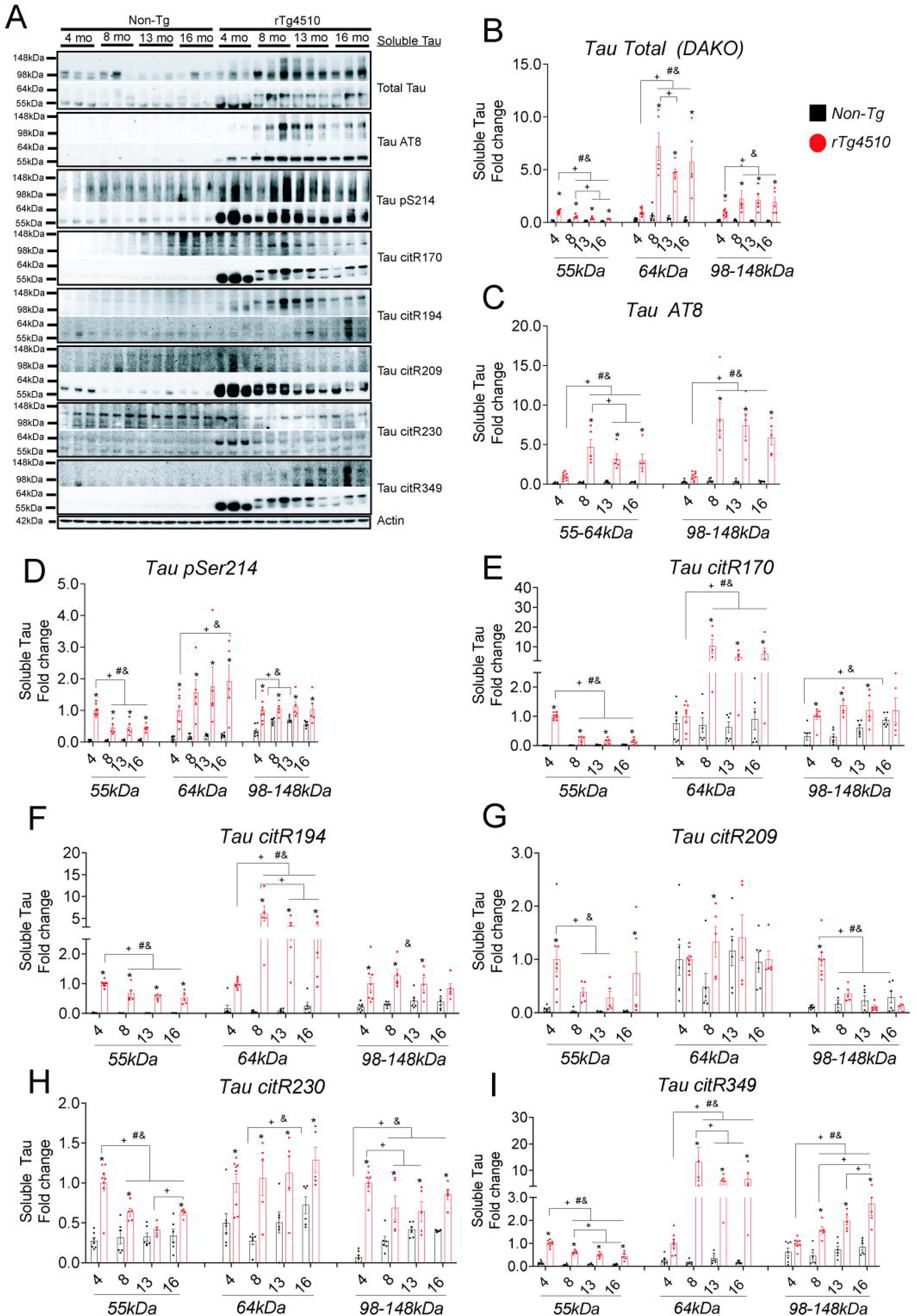
Age analysis of soluble tau progression in rTg4510 mice. Western blot analysis of cortical brain homogenate from soluble tau fraction in 4, 8, 13, 16 months of rTg4510 mice and age matched non-transgenic littermates probed with total tau (DAKO) (**A, B**), tau AT8 (**A**, **C**), tau pS214 (**A**, **D**), tau citR170 (**A**, **E**), tau citR194 (**A**, **F**), tau citR209 (**A**, **G**), tau citR230 (**A**, **H**), and tau citR349 (**A**, **I**) antibodies at molecular weights 55kDa, 64kDa, and high molecular weight tau (98-148kDa). Values normalized to actin and 4 mo-old rTg4510 intensity were presented as fold change. Statistical analysis were performed by two-way ANOVA; &=denotes significance by genotype, # =by age, * =pairwise comparison between genotype within the age group, ^+^ =denotes pairwise comparison of rTg4510 across age P < 0.05 (n=3-8/ age).

Next we measured detergent sarkosyl soluble tau in mice of the same age. Overall, there was a main effect of age and genotype for all epitope species measured between rTg4510 mice and non-transgenic littermates. Several epitopes from rTg4510 mice displayed a similar pattern to that of the soluble tau. Specifically, total tau (**Figure 10A, B**), tau pSer214 (**Figure 10A, D**), tau citR209 (**Figure 10A, F**), tau citR194 (**Figure 10A, G**), tau citR230 (**Figure 10A, H**) (55kDa) increased at 4-months, then subsequently decreased between 8-16-months. Tau citR349 increased at 4-months and remained elevated until 13-months (**Figure 10A, I**). Sarkosyl soluble (64kDa) total tau (**Figure 10A, B**), tau pSer214 (**Figure 10A, D**), and tau AT8 (**Figure 10A, C**) increased at 4-months and further accumulated at 8-months but also remained elevated. Interestingly, sarkosyl soluble (64kDa), tau citR170 (**Figure 10A, E**), tau citR194 (**Figure 10A, G**), tau citR230 (**Figure 10A, H**), and tau citR349 (**Figure 10A, I**) all increased at 8-months but subsequently dropped thereafter. Very little tau citR170 was detected in the sarkosyl soluble fraction apart from a 64kDa band at 8-months (**Figure 10A, E**). Tau AT8 (55-64kDa) slightly increased at 4-months and further accumulated at 8-16 months (**Figure 10A, C**). We detected increased high molecular tau multimers (98-148kDa) for total tau (**Figure 10A, B**), AT8 (**Figure 10A, C**), tau pSer214 (**Figure 10A, D**) at 8-months which remained elevated throughout the study however citR tau multimers were not readily detectable.

**Figure 10.**
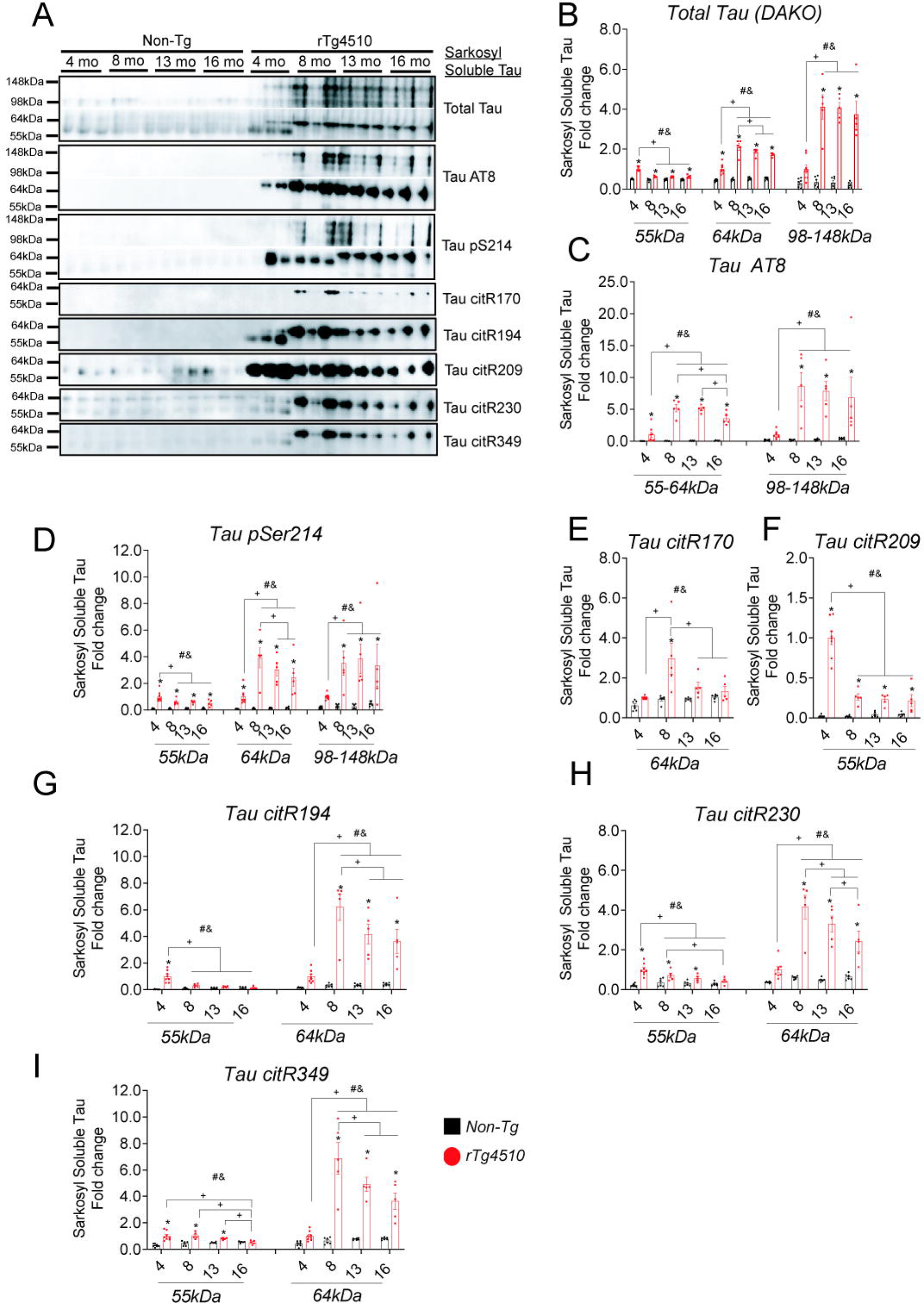
Age analysis of sarkosyl soluble tau progression in rTg4510 mice. Western blot analysis of cortical brain homogenate from sarkosyl soluble tau fraction in 4, 8, 13, 16 months of rTg4510 mice and age matched non-transgenic littermates probed with total tau (DAKO) (**A, B**), tau AT8 (**A**, **C**), tau pS214 (**A**, **D**), tau citR170 (**A**, **E**), tau citR209 (**A**, **F**), tau citR194 (**A**, **G**), tau citR230 (**A**, **H**), and tau citR349 (**A**, **I**) antibodies at molecular weights 55kDa, 64kDa, and high molecular weight tau (98-148kDa). Values normalized to actin and 4 mo-old rTg4510 intensity were presented as fold change. Statistical analysis were performed by two-way ANOVA; &=denotes significance by genotype, # =by age, * =pairwise comparison between genotype within the age group, ^+^ =denotes pairwise comparison of rTg4510 across age P < 0.05 (n=3-8/ age).

Finally, we measured sarkosyl insoluble tau in rTg4510 mice and non-transgenic littermates (**Figure 11**). Overall, there was a main effect of age and genotype for total tau, AT8, tau pSer214, tau citR194, and a genotype affect for tau citR170 and tau citR349. Although no total tau (**Figure 11A, B**), AT8 (**Figure 11A, C**), or taupSer214 (**Figure 11A, D**) was detected at 4-months of age the 55kDa-64kDa and high molecular weight (98-148kDa) increased at 8-months and remained elevated to 16-months (**Figure 11**). Tau citR170 (**Figure 11A, E)**, tau citR194 (**Figure 11A, F**), and tau citR 349 (**Figure 11A, G**) was only detectable after >2-3 hour manual exposure in several mice. However, only a 64kDa band of tau citR170 was detected at 8-months and remained elevated at 8-16 months. Tau citR194 (55kDa) increased 8-16 months, while the high molecular weight (98-148kDa) increased at 8- and 16 months. Tau citR349 (55kDa and 98-148kDa) also increased at 8- and 13-months but waned at 16-months. These data showed that both TBS-soluble and sarkosyl soluble (55kDa) phospho tau pSer214 and citR tau species increase early during tau pathology but subsequently decreased and remained low, while accumulating a 64kDa band in both fractions. Conversely, tau AT8 (64kDa and 98-148kDa) accumulated later in disease and remained elevated throughout the duration of the study. Importantly, sarkosyl insoluble tau readily accumulated phospho tau AT8 and total tau however citrullinated tau was not as readily detected until after 2-3hrs of manual exposures. This suggest that the majority of citR tau species does not particularly accumulate in the insoluble fractions like AT8. These data could also suggest an overall lower abundance of citR tau compared to phospho tau species, however further research is necessary to understand the solubility profile of citR tau.

**Figure 11.**
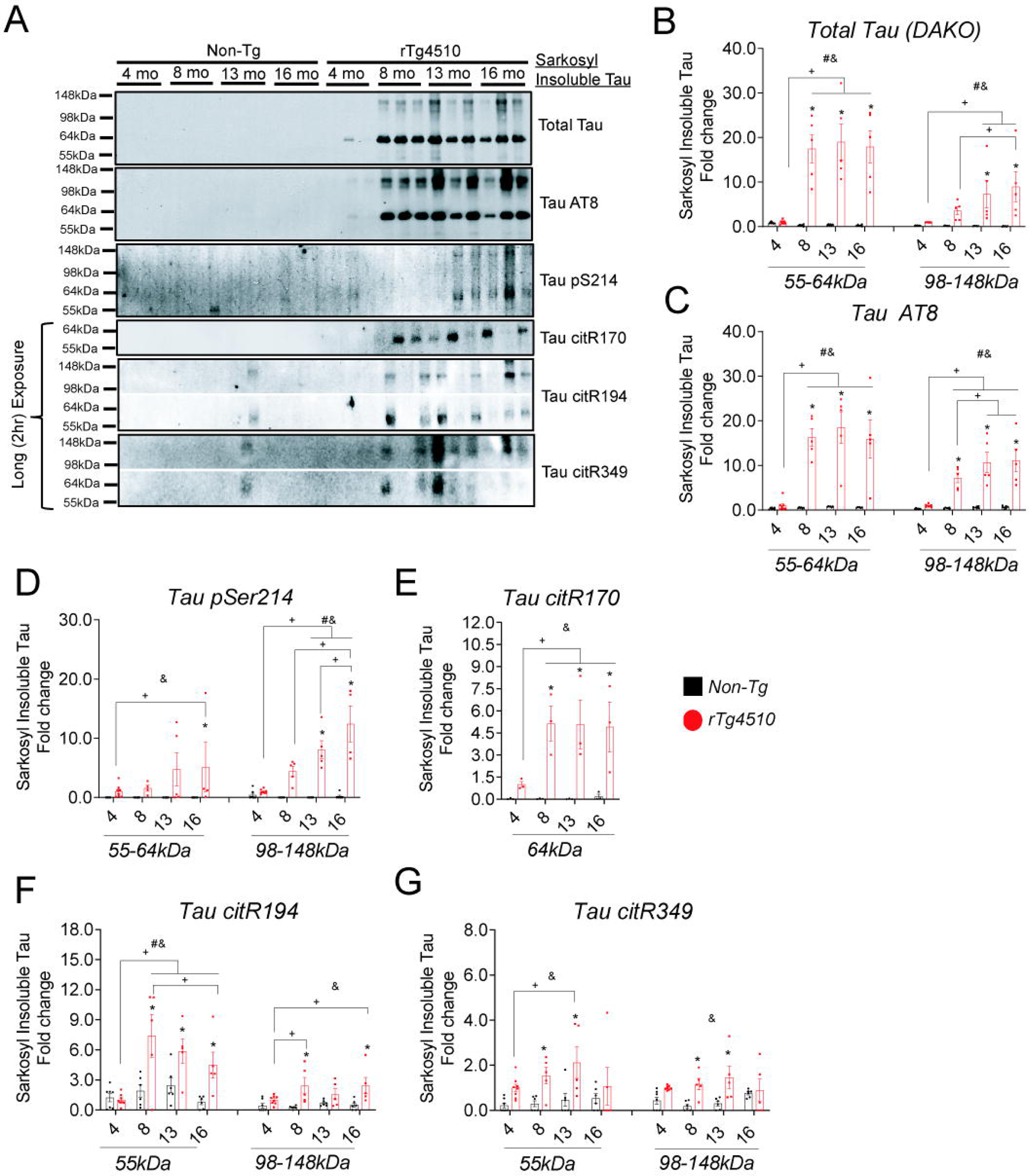
Age analysis of sarkosyl insoluble tau progression in rTg4510 mice. Western blot analysis of cortical brain homogenate from sarkosyl insoluble tau fraction in 4, 8, 13, 16 months of rTg4510 mice and age matched non-transgenic littermates probed with total tau (DAKO) (**A, B**), tau AT8 (**A**, **C**), tau pS214 (**A**, **D**), tau citR170 (**A**, **E**), tau citR194 (**A**, **F**), tau citR349 (**A**, **G**) antibodies at molecular weights 55kDa, 64kDa, and high molecular weight tau (98-148kDa). Values normalized to actin and 4 mo-old rTg4510 intensity were presented as fold change. Statistical analysis were performed by two-way ANOVA; &=denotes significance by genotype, # =by age, * =pairwise comparison between genotype within the age group, ^+^ =denotes pairwise comparison of rTg4510 across age P < 0.05 (n=3-8/ age).

### Tau citrullination in Alzheimer’s disease brains

Next, we measure citrullinated tau (tau citR170, citR194, citR209, citR230, citR349) in the same cases positive for phospho tau (**Figure 1**). No positive staining of citR tau was observed in controls with Braak II (**Figure 12A**). Occasionally, we observed rare staining of neuronal tau citR194, citR209, citR230, and citR349 in the Braak IV case (**Figure 12A**). However, tau citR170, citR194, citR209, citR230, citR349 increased in Braak VI cases and consisted of tangle-like pathology, neuropil threads, and neuritic plaques (**Figure 12A**). Similar to phospho tau, we performed western blot analysis and found that tau citR170, tau citR194, tau citR230, and tau citR349 (**Figure 12B, C**) increased in AD (+)/ AT8 (+) samples. Interestingly, tau citR209 monomer decreased in western samples in AD+ tissue. This cause of the discrepancy between immunohistochemisty and western blot remains unclear but several possible explanations should be considered. The tau citR209 antibody could bind with higher affinity to tau and recognize structurally distinct proteoforms by immunohistochemistry that are denatured by SDS-PAGE. Another possible explanation may reflect the epitope stretch itself. The citR tau antibody specifically recognizes the citrullinated moiety of R209, however there are several potential post-translational modifications including citrullination and phosphorylation surrounding the epitope sequence including *R^209^SR^211^T^212^PS^214^ LPT^217^PPTR^221^*. We show that citR209 is citrullinated but also demonstrate that tau citR211 (by mass spectrometry) and tau citR221 (by mass spectrometry and animal models) can also possibly be citrullinated. In addition, this area is heavily phosphorylated early in disease including tau pThr212, tau pSer214, tau pThr217 and serves as early AD biomarkers (Ashton, Brum et al. 2024). It remains possible that early phosphorylation of these sites reduces citrullination of surrounding epitopes or precludes antibody binding to the citR209 site under western blotting conditions. It should be noted that tau pSer214 increased in both AD (+)/ AT8 (−) or AD (+)/ AT8 (+), whereas tau citR209 decreased in both AD (+)/ AT8 (−) or AD (+)/ AT8 (+), however this pattern was not observed at other citR tau sites. Nonetheless, further research is warranted to determine exact interactions around this epitope sequence.

**Figure 12.**
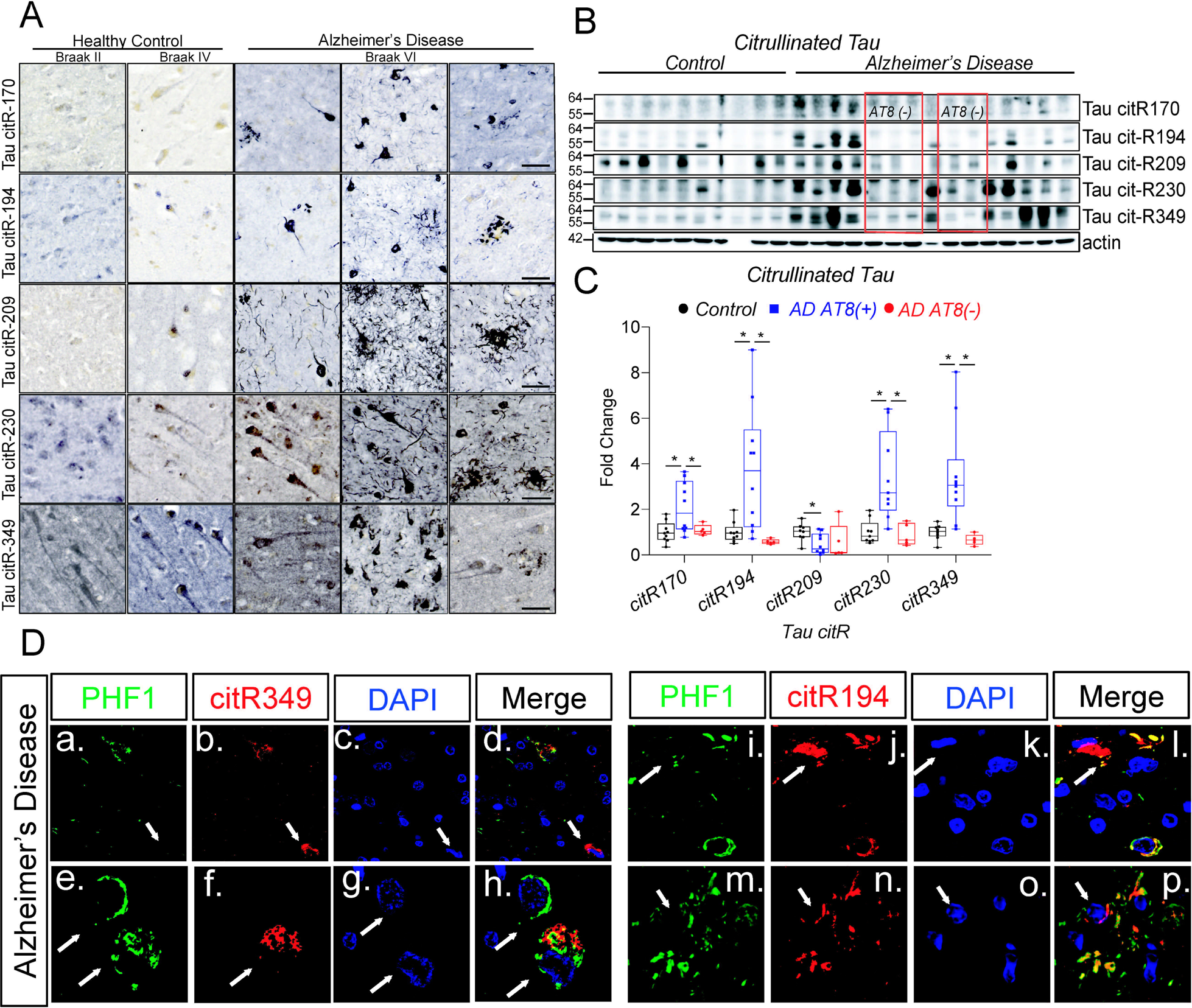
Citrullinated tau is increased in Alzheimer’s disease brains. Panel **A** shows immunohistochemical staining of tau citR170, R194, R209, R230, R349 in healthy controls (Braak II), Braak IV, and Alzheimer’s disease (AD) Braak VI (**A**). Panel **B** shows western blot analysis of cortical homogenates from AD brain and unaffected controls and densitometry analysis (**C**) of tau citR170, R194, citR209, R230, R349 in AT8 (+) AD samples compared to AT8(−) AD and unaffected control samples. Raw values were normalized to actin and graphed as fold change (± SEM). Statistical analysis performed by One-way Anova with Fisher’s LSD as pos hoc (*P < 0.05, n=10/ control sample, n=10/ AD AT8 (+), n=5 AD AT8 (−)). Panel **D** shows double labeling of phospho tau PHF1 (green), tau citR349 (red), nuclei (DAPI-blue) (**D. a-h**) and phospho tau PHF1 (green), tau citR194(red), nuclei (DAPI-blue) (**D. i-p**). Images show that PHF1 labels neurons independent and together with tau citR349 (**D. a-h**, arrows) and tau citR194 (**D. i-p**) but may also locate in different subcellular compartments of the neuron. Additionally, tau citR349 (**D. a-d,** arrow) and tau citR194 (**D. i-l**, arrow) label neurons independent of phospho tau PHF1.

Finally, given that citR tau expression associated with phospho tau in AD brains, we double labeled PHF1 with two citR tau epitopes including tau citR349 and tau citR194 (**Figure 12 D**). We found that PHF1 co-labeled neurons, plaques, and neuropil threads with both tau citR349 and tau citR194 but also labeled cells negative for citR tau. Conversely, tau citR349 and tau citR194 labeled neurons that were PHF1 negative indicating that both pathways can occur simultaneously and independently. Additionally, some cells were positive for both PHF1 and citR tau but failed to co-localize in the same region or overlap possibly suggesting different activation profiles, subcellular compartmentalization, or different proteoforms of tau.

## Discussion

In the current study, we identified citrullination of tau on multiple epitopes under several conditions including recombinant protein reactions, cellular models, animal models of tauopathy, and human tissue. Several reports demonstrated increased PAD2/PAD4 activity and hypercitrullination in Alzheimer’s disease brain (Ishigami, Ohsawa et al. 2005). We also confirmed those reports in a small cohort of AD and control samples, but similarly showed that PAD2 and PAD4 expression increased in brains with higher phospho tau burden. Although PAD2 and PAD4 protein expression increased in AD, protein abundance may not be a rate limiting step to increase citrullination activity. PADs are seemingly coordinately activated with calcium load. Recent evidence suggests five different AD subtypes that distinguish molecular processes including neuronal hyperplasticity, innate immune activation, RNA dysregulation, choroid plexus dysfunction, and blood brain barrier impairment (Tijms, Vromen et al. 2024). Neurons with increased activity secreted more amyloid beta and tau were associated with the hyperplasticity subtype. Further work is warranted to understand if neuronal hyperplasticity subtypes are associated with aberrant calcium dysfunction, hypercitrullination and thereby hyper-citrullinated tau.

We also demonstrated that tau could be fully citrullinated using recombinant protein systems. Full-length tau (2N4R) can become citrullinated by PAD4 or PAD2 at all 14 arginine residues with recombinant protein during optimal buffer conditions; however it remains unclear if all tau isoforms or strains are citrullinated to the same extent *in vivo*. It is not entirely clear how citrullination of tau impacts its function, metabolism and aggregation profile but is indeed necessary to uncover given that citrullination may seemingly be irreversible. We simulated pseudo-citrullination on tau with arginine to glutamine residues to begin to understand the disorder ranking of the modification. This change in modification was not an obvious conclusion based on changes in the charge neutralization, increased hydrophobicity, and reduced π-π interactions. Increased hydrophobicity should typically decrease disorder potential, whereas increased mean net charge, which is the case for tau citrullination associates with increased disorder propensity. Altogether, computational analysis suggests that because of the increased local hydrophobicity, citrullinated tau might be slightly more ordered, form spontaneous LLPS, and prone to more aggregation due to the increased potential to form oligomers. However, experimental confirmation is needed to understand the full extent of citrullinated tau’s biophysical properties. Not only does arginine residues drive salt bridge formation with glutamate and aspartate residues (Walker and Causgrove 2009, Meuzelaar, Vreede et al. 2016) and promote alpha helix H-bond stability, a recent report showed that arginine π-π□bonds within tau were key determinants for tau’s protein network and demonstrated how arginine’s drive aberrant interactions of proteins with tau fibrils (Ferrari, Stucchi et al. 2020). The higher-order aggregation from monomer to fibril formation rewires avidity properties of tau and attracts new sets of interactor proteins for the tau proteome. In contrast, the citrulline moiety promotes protein unfolding, increases hydrophobicity, reduces the overall net charge, causes a loss of ionic bonds, and interference with H-bond stabilization (Tarcsa, Marekov et al. 1996). The same report demonstrated that the degree and rate of arginine to citrulline conversion by PAD directly correlated with the structural order of the substrate proteins (i.e., filaggrin, trichohyalin). The authors illustrated that filaggrin, a disorder protein, quickly underwent up to 95% citrulline conversion. However, citrullination proceeded more slowly to about 25% with another protein, trichohyalin, which is highly enriched in α-helix moieties.

Although increased citrullination associates with AD, Prion disease (Jang, Kim et al. 2008), Multiple sclerosis (Moscarello, Mastronardi et al. 2007, Witalison, Thompson et al. 2015), causal outcomes of aberrant PAD activity still remain unclear (Ishigami, Ohsawa et al. 2005, Acharya, Nagele et al. 2012, Pietronigro, Della Bianca et al. 2017). We focused on PAD4 induced tau citrullination primarily due to its neuronal expression in the brain. Given that PAD2 has been reported to primarily express in astrocytes and glial also observed in this study, it is conceivable that PAD2 expression in rTg4510 mice increased in glia secondarily to the tau endophenotype, however more work is needed to validate the specific role of PAD2 during tau deposition. This also raises the immediate question of astrocytic tau in tauopathies like Age-Related Tau Astrogliopathy (ARTAG), Progressive supranuclear palsy (PSP). Tau induction in iHEK tau cells also increased PAD4 but not PAD2 suggesting a bidirectional relationship between tau and PAD4 at the molecular level. Interestingly, tau citrullination occurred early in rTg4510 mice, which tracked with early PAD4 expression in mice. PAD4 harbors SNPs that associates with AD risk (*rs16824888*) (Nazarian, Arbeev et al. 2019) and amyotrophic lateral sclerosis (ALS) (*rs2240335*) (Tanikawa, Ueda et al. 2018) and loss or gain of function may differentially promote or subvert specific proteinopathies. One report (Tanikawa, Ueda et al. 2018) described how PAD4 impacted protein aggregation and susceptibility to ALS. The authors identified 159 substrates in cells (tau was not among those identified), with an RG/ RGG motif preference for PAD4 and suggested that citrullination increased solubility of FET proteins to reduce their aggregation following stress paradigms. PAD4 induced citrullination prevented methylation and aggregation through stress granules of FET proteins (**<u>F</u>**US, **<u>E</u>**WS, **<u>T</u>**AF15). This is an interesting result and might suggest a alternative interpretation based on our computer analysis of “pseudo-citrullinated tau”. Tau and by extension tauopathies are unique in that soluble aggregates, insoluble aggregates, different proteoforms and tangles all associate with some measure of toxicity but manifest diverse clinical syndromes, variable phenotypes, and regionally accumulate in different areas of the brain depending on the tau variant or strain. Increasing evidence suggest that tau forms soluble aggregates (soluble oligomers) versus folded (fibrillar oligomers); however it is unclear how these proteoforms emerge and which species is more toxic (Ghag, Bhatt et al. 2018). It is also plausible that irreversible citrullination locks tau into certain proteoforms or oligomeric species. Tau contains two RG/ RGG motifs, one at tau R^155^G and the other at tau R^194^XG, but can also be citrullinated at multiple sites that do not contain these motifs suggesting that other mechanisms that govern PAD induced citrullination of tau.

Although many of the citrullinated tau epitopes were not yet tested or shown for AD tissue (tau citR170, tau citR194, tau citR209, tau citR231, tau citR349 shown) in the current study, 10 citR tau antibodies labeled tau in mouse models of tauopathy (shown: tau citR170, tau citR194, tau citR209, tau citR231, and tau citR349; not shown; tau citR5, tau citR23, tau citR155, tau citR221, tau citR379, tau citR406). We sought out to create an antibody panel to test new hypotheses for domain-specific functions of tau but to also examine how PAD’s citrullinate various epitopes on tau. One particular question was how calcium load impacts PAD4’s substrate specificity on various tau epitopes. We found that higher concentrations of PAD4 (>500nM) citrullinated several arginine’s on tau even in the absence of calcium, whereas low concentrations of PAD4 (approximately 10-100 fold less) citrullinated specific arginine residues on tau only in the presence of high calcium (10mM). These data suggest that low abundance PAD4 with calcium modifies the specificity of citrullinated substrates on tau. It also suggests that aberrant expression of PAD4 potentially with or without calcium causes low substrate specificity possibly leading to hypercitrullination. Of note, these conclusions are in context with recombinant protein systems under optimal buffer conditions. However, our data also provide some context as to the versatility of PAD-induced citrullination and depends on calcium load, PAD levels, and the substrate/ epitope client itself. We induce rapid tau expression with tetracycline in iHEK Tau cells and showed that PAD4 also increased in response to tau suggesting a possible feed forward cycle of hypercitrullination. From a therapeutic angle, potency and efficacy of PAD inhibitors will need to be tailored to disease conditions based on these considerations.

Our data also indicated that PAD4 citrullinates tau at arginine residues within the N-terminal, proline rich region, microtubule binding domain, and c-terminal domain, suggesting broad changes in functional activity of tau. Interestingly, the highest stretch of arginine residues and citrullinated sites occurs within the proline rich region. This domain may serve as a scaffold and signaling hub as previously suggested (Mueller, Combs et al. 2021). However, it is not clear how arginine’s influence these interactions. Recent development of PAD inhibitors has made hyper citrullination a therapeutic target in animal models of ischemic stroke (Kim, Lee et al. 2019), multiple sclerosis (Moscarello, Lei et al. 2013, Tejeda, Bello et al. 2017), TBI (Vaibhav, Braun et al. 2020), spinal cord injury (Feng, Min et al. 2021), neonatal hypoxic ischemia (Lange, Rocha-Ferreira et al. 2014). Here, we also validated that PAD inhibitor BB-CL-amidine prevented citrullination of recombinant tau. Although we do not know how citrullinated tau impacts neuronal toxicity, future studies will be important to test PAD inhibitors in models of tauopathies.

In conclusion, we identified that tau can become fully citrullinated at all 14 arginine residues to some extent through PAD4 and PAD2 and that novel antibodies label citR tau in animal models of tauopathy and AD brains. Our findings indicate that citrullination of tau occurs in AD neuropathologic change. Future studies will be important to understand the full extent of tau citrullination in other tauopathies and the functional impact of this post-translational modification on tau. Additionally, identifying the regulatory mechanisms of PADs during AD pathogenesis and tauopathies will also be important therapeutically. Together, this work provides a new area of tau biology that warrants further understanding and consideration as a potential target for anti-tau approaches.

## Acknowledgment

We would like to thank Dr. Rakez Kayed for his generous gift of anti-oligomeric tau (T22). We also thank the NIH NeuroBioBank donors and their families for their generosity.

## Funding

DL-1RF1AG072728; DL-Brightfocus Foundation; DL-University of Kentucky Neuroscience Research Priority Award (NRPA); MLBS-R01NS123454; MLBS-CART Rotary; P30 AG028383

## Conflict of interest

The authors have no conflict of interest.

## Abbreviations

PAD: peptidyl arginine deiminase
PTM: post-translational modification
CitR: Citrullination/ citrullinated
AD: Alzheimer’s disease
ALS: Amyotrophic lateral sclerosis
FTD: frontotemporal dementia

## Supplementary Figures/Tables

**Supplementary Table 1.** List of citrullinated tau epitopes by PADs as identified by Mass spectrometry analysis. Localization probability =1 present probability of occurrence, and digested peptide intensity is shown.

**Supplementary Figure 1.** Mass spectrometry spectra of arginine citrullination sites on tau protein following incubation with PAD2 and PAD4 as identified by mass spectrometry. Also shown are spectra of citR tau and retention times.

## Notes

### Competing Interest Statement

The authors have declared no competing interest.

