## Supplementary figures and images for "Probing tau citrullination in Alzheimer’s disease brains and mouse models of tauopathy"

### Supplemental Figure 1

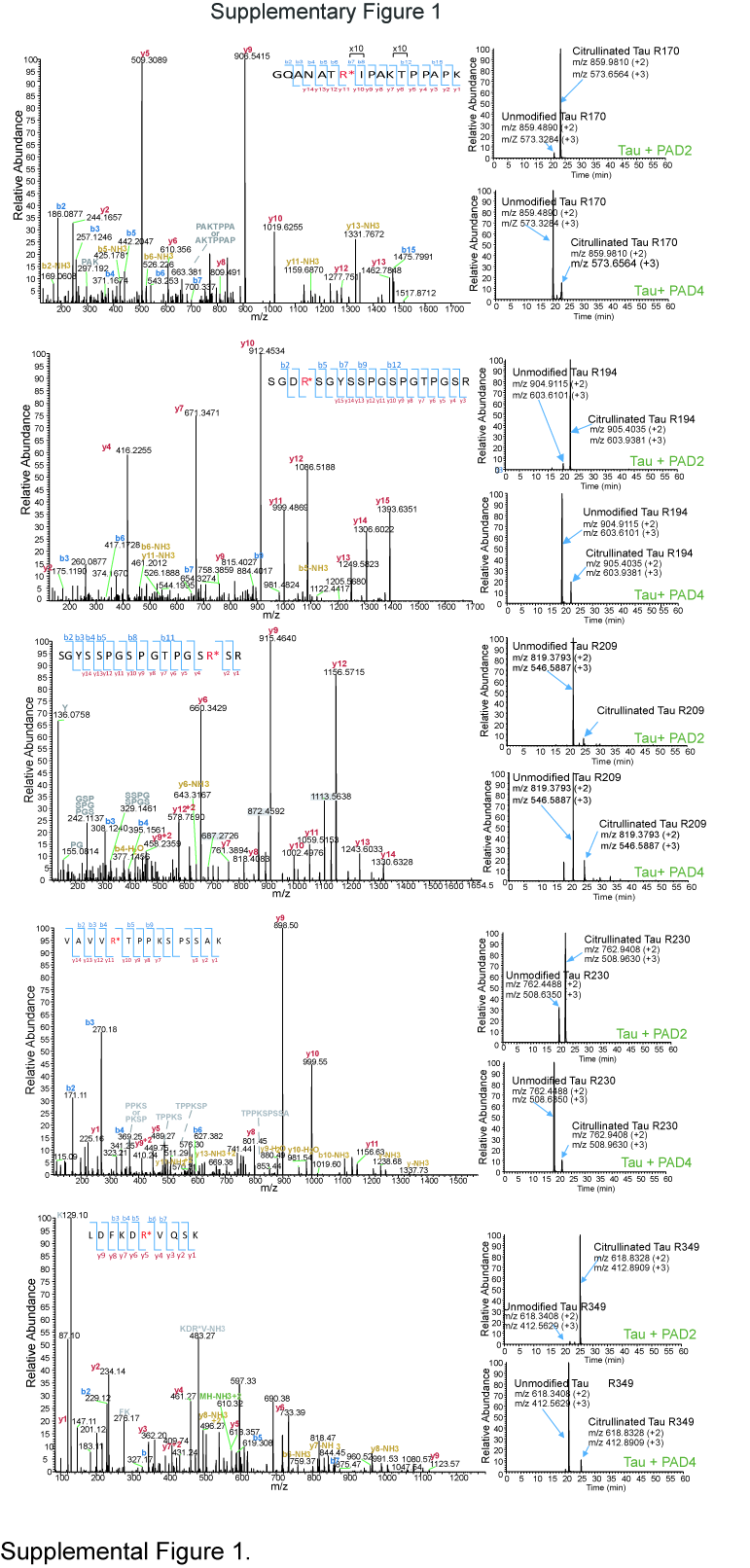

### Supplemental Table1

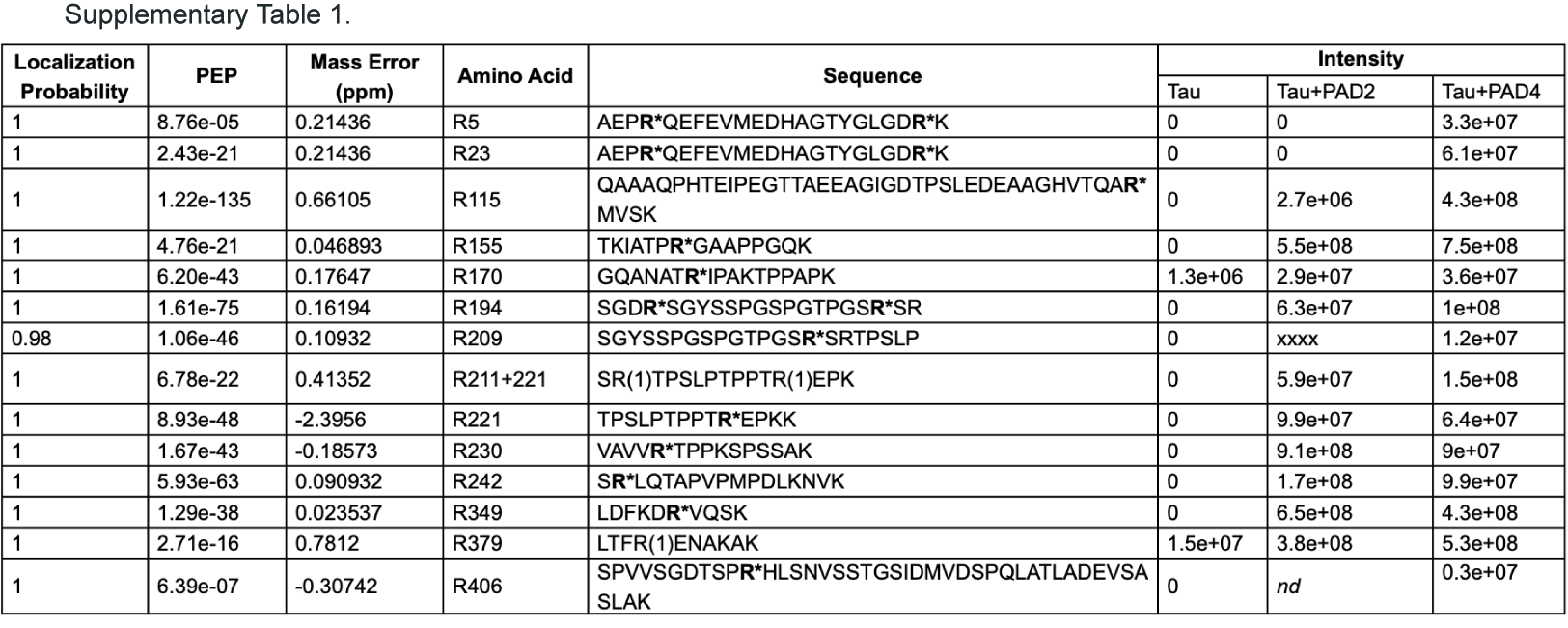
